# Targeted hybridization capture enables comprehensive detection of freshwater bioassessment invertebrates from environmental DNA

**DOI:** 10.64898/2026.08.14.744904

**Authors:** Joseph M. Craine, John L. Darcy, Jessica Devitt, Devin Leopold, Galen W. Miller, Mitchell Ralson, Nick Schulte, Noah Fierer

## Abstract

Freshwater bioassessment relies on assessing aquatic assemblages to infer ecological conditions, yet conventional surveys require extensive field sampling, specimen processing, and specialized taxonomic expertise. Existing environmental DNA (eDNA) methods have not yet provided a practical alternative to conventional macroinvertebrate assays in part because current approaches cannot feasibly recover broad taxonomic diversity at sufficient taxonomic resolution. Here, we evaluated targeted hybridization capture of mitochondrial cytochrome oxidase I (COI) target sequences as a unified molecular approach for cross-phylum freshwater bioassessment. Environmental DNA was collected at 18 sites along 63 km of Boulder Creek spanning nearly 1,500 m of elevation from forested headwaters to agricultural plains. COI targets were enriched using custom RNA bait panels designed to target regional freshwater arthropods, annelids, and molluscs. Hybridization capture increased recovery of COI sequences ∼1,760-fold relative to unenriched shotgun libraries, generating Folmer-region COI contigs that averaged ∼400 bp. Across the watershed, we recovered sequences for approximately 450 macroinvertebrate genera across 8 phyla. Detected macroinvertebrate richness averaged 56 genera per site and increased down Boulder Canyon before declining downstream of the city. Macroinvertebrate assemblage composition from hybridization capture paralleled patterns observed with past conventional bioassessment. These results demonstrate that targeted hybridization capture enables robust, cross-phylum detection of species used for freshwater bioassessment from environmental DNA.

## Introduction

Freshwater ecosystems provide essential services including biodiversity conservation, water purification, nutrient cycling, fisheries production, and recreation, yet they are among the most anthropogenically altered ecosystems worldwide (Dudgeon et al. 2006; Reid et al. 2019). Protecting these systems across broad spatial scales requires routine assessment of ecological condition to detect impairment so that stressors can be relieved and condition restored. Because aquatic organisms integrate environmental conditions over weeks to years and respond predictably to a broad range of physical, chemical, and biological stressors, biological communities can provide more comprehensive measures of ecological health than instantaneous physicochemical measurements alone (Karr 1981; Barbour et al. 1999; Bonada et al. 2006). Consequently, freshwater biomonitoring programs routinely evaluate assemblages of fish, aquatic macroinvertebrates, diatoms, bacteria, or amphibians using multimetric indices that have become the foundation of water quality assessment and environmental management worldwide (Karr 1981; Hering et al. 2006; Vadas et al. 2022).

Despite their importance, conventional biological surveys remain labor-intensive, expensive, and dependent upon specialized taxonomic expertise (Bonada et al. 2006). Standard assessments typically require extensive field sampling, with separate efforts for each target group, e.g., electrofishing or seines for fish, kick nets or Surber samplers for macroinvertebrates, rock scrubbing or scraping for periphyton diatoms. Depending on the target group, manual sorting and morphological identification can often require days to weeks of subsequent effort (Barbour et al. 1999; Bonada et al. 2006). Many taxa, particularly aquatic insects, can only be reliably visually identified to genus or family because diagnostic morphological characters are absent or require life stages that are rarely collected (Grover et al. 2022; Merritt, Cummins, and Berg 2019). Moreover, skilled taxonomists are increasingly too rare to process samples at the scale needed for broad-scale monitoring. These constraints limit the spatial and temporal resolution of monitoring programs, their effectiveness in identifying stressors, and contribute substantially to program costs. Improving the speed, taxonomic resolution, and cost-efficiency of aquatic biomonitoring would allow expansion of assessment efforts, leading to better characterization of ecological condition at individual sites and across regions. DNA metabarcoding of bulk organism collections has streamlined taxonomic identification for many applications, but it still relies on collecting organisms from the ecosystem, generally requires sorting and processing bulk samples, and does not eliminate the logistical burden of field collections (Elbrecht and Leese 2017; Porter and Hajibabaei 2018). Environmental DNA (eDNA) has emerged as a promising alternative because organisms can be detected directly from DNA released into water (Taberlet et al. 2012; Deiner et al. 2017). With eDNA analysis, a single water sample can simultaneously characterize multiple taxonomic groups and can be processed using highly standardized laboratory workflows (Deiner et al. 2017; Taberlet et al. 2018). Over the past decade, eDNA has transformed biodiversity surveys for invasive species detection, endangered species monitoring, community ecology, and increasingly, freshwater bioassessment (Deiner et al. 2017; Altermatt et al. 2025).

Despite the promise of aquatic eDNA analyses, no current molecular approach using aquatic eDNA fully satisfies the requirements of comprehensive aquatic biomonitoring. Species-specific quantitative PCR assays provide high sensitivity but are inherently limited to a small number of target taxa and can suffer from false positives without secondary sequencing of amplification products (Goldberg et al. 2016). Shotgun metagenomic sequencing avoids amplification bias but typically requires prohibitively deep sequencing because informative DNA regions from target organisms represent only a minute fraction of total environmental DNA (Deiner et al. 2017; Bohmann et al. 2014). Metabarcoding has become the dominant approach for assemblage-level eDNA analysis because it characterizes broad sets of species (Deiner et al. 2017; Taberlet et al. 2018). For freshwater macroinvertebrates, most eDNA studies target mitochondrial cytochrome oxidase I (COI), which provides the most comprehensive reference databases and high species-level taxonomic resolution available for metazoans (Hebert et al. 2003; Ratnasingham and Hebert 2007). However, comprehensive recovery of macroinvertebrate diversity from eDNA remains constrained by an inherent tradeoff in PCR primer design. COI-based primers that maximize taxonomic coverage typically co-amplify abundant non-target DNA, including oomycetes, diatoms, protists, and other eukaryotes, substantially reducing sequencing efficiency and recovery of target macroinvertebrates (Hajibabaei et al. 2019; Macher et al. 2018; Leese et al. 2021; Fueyo et al. 2025). Reducing non-target amplification through primer design increases the proportion of target sequences in the final library, but this improved specificity is coupled with reduced detection of many desired taxa, such as flatworms, molluscs, isopods, and key insect genera (Leese et al. 2021). Consequently, no single COI primer pair efficiently amplifies the full phylogenetic breadth of target freshwater macroinvertebrates while simultaneously reducing amplification of abundant non-target DNA. Although existing eDNA metabarcoding assays have demonstrated sufficient taxonomic coverage to generate bioassessment metrics that recover some aspects of ecological condition across freshwater systems (Pawlowski et al. 2018; Hering et al. 2018), they are unlikely to achieve comprehensive recovery of freshwater diversity from environmental DNA using a single metabarcoding assay, or even a small number of assays.

Targeted hybridization capture offers a fundamentally different approach to target sequence enrichment, combining the taxonomic breadth of shotgun sequencing with the sequencing efficiency of targeted amplicons (Mamanova et al. 2010; Horn 2012). Rather than using PCR primers to selectively amplify target molecules, targeted capture enriches sequencing libraries through sequence-specific hybridization. Sets of biotinylated RNA molecules (“baits”) are constructed to match portions of target sequences and then allowed to hybridize to homologous and near-homologous DNA fragments from the total pool of eDNA. With this approach, target molecules can be isolated from non-target DNA prior to sequencing. Because enrichment is achieved through thousands of independently designed hybridization probes rather than a single pair of universal primers, target specificity and taxonomic breadth can be optimized simultaneously. In addition, hybridization tolerates sequence divergence, allowing recovery of homologous loci across broad phylogenetic groups and genomic regions. Because targeted capture uses an array of probes that tile across a target region, it can recover overlapping DNA fragments that assemble into contigs longer and more complete than PCR-based metabarcoding amplicons. Targeted capture has become a widely adopted tool in phylogenomics for recovering conserved nuclear loci across divergent taxa (Faircloth et al. 2012; Lemmon, Emme, and Lemmon 2013) and in ancient DNA research by enabling efficient enrichment of degraded DNA from archaeological and paleoecological specimens (Carpenter et al. 2013; Paijmans, González Fortes, and Förster 2019).

Despite these advantages, targeted capture has not yet been widely adopted for freshwater eDNA bioassessment. Targeted capture has only occasionally been applied to environmental DNA. Isolated studies have included a freshwater biodiversity survey with limited taxonomic richness (Wilcox et al. 2018), terrestrial vertebrate detection in African and Cambodian waterholes (Seeber et al. 2019; Li et al. 2023), and recovery of whale shark mitochondrial and nuclear DNA from seawater (Jensen et al. 2021). Collectively, these studies demonstrate that target enrichment can recover informative genetic sequences from complex environmental samples, but have yet to show the technique can be used to recover diversity suitable for bioassessment.

Technical challenges remain across multiple stages of the targeted capture workflow. Designing comprehensive bait panels requires balancing taxonomic breadth with probe redundancy while accommodating extensive sequence diversity and incomplete reference databases. Laboratory protocols for capture efficiency, library preparation, and sequencing need to be optimized and are less standardized than PCR-based approaches (Horn 2012). Bioinformatic analysis also presents unique challenges, including assembly of enriched mitochondrial sequences, accurate taxonomic assignment across diverse taxa, and integration of recovered communities into ecological analyses.

Here, we evaluate whether targeted capture can provide a practical framework for comprehensive, cross-phylum detection of species used in freshwater bioassessment from a single environmental DNA workflow. We developed custom RNA bait panels comprising 12,000 individual bait sequences that target mitochondrial COI sequences from freshwater macroinvertebrates species. These panels were applied with hybridization reactions to environmental DNA collected from 18 sites spanning approximately 60 km of Boulder Creek, Colorado. The sampling transect encompassed nearly 1,500 m of elevation change, extending from high-elevation conifer forests through montane canyon, urban, and downstream agricultural ecosystems. We evaluated target enrichment efficiency, assembly performance, taxonomic recovery, and longitudinal biodiversity patterns, as well as a few established bioassessment indices directly from the capture data. To assess ecological validity, we compared the recovered assemblages with morphological data from a long-term state monitoring program. Together, these analyses assessed whether targeted capture can overcome key limitations of existing eDNA approaches while producing ecologically meaningful characterization of assemblages across diverse freshwater taxonomic groups.

## Materials and Methods

### Study area

The study was conducted in the Boulder Creek watershed, Colorado, USA, spanning approximately 65 km from the headwaters near Lost Lake (39.95390, –105.61980) to the confluence of Boulder Creek with the St. Vrain River (40.17042, –104.99156). The sampling transect encompassed nearly 1,500 m of elevation (2,943–1,475 m) and traversed four major watershed reaches representing distinct ecological and land-use zones: Upper Canyon, Lower Canyon, Boulder urban, and Plains Reach (**Figure 1**). Sites extended from high-elevation conifer forest through montane canyon, urban Boulder, and downstream agricultural landscapes. A total of 18 sampling sites were established along the watershed (**Figure 1**, **Table S1**). All field collections were completed during a single day (May 27, 2026) to minimize temporal variation in assemblage composition. The USGS gaging station for Boulder Creek at North 75^th^ St. (USGS-06730200) on that day registered a depth of 1.55 m and a discharge of 2.66 m^3^s^-1^.

**Figure 1.**
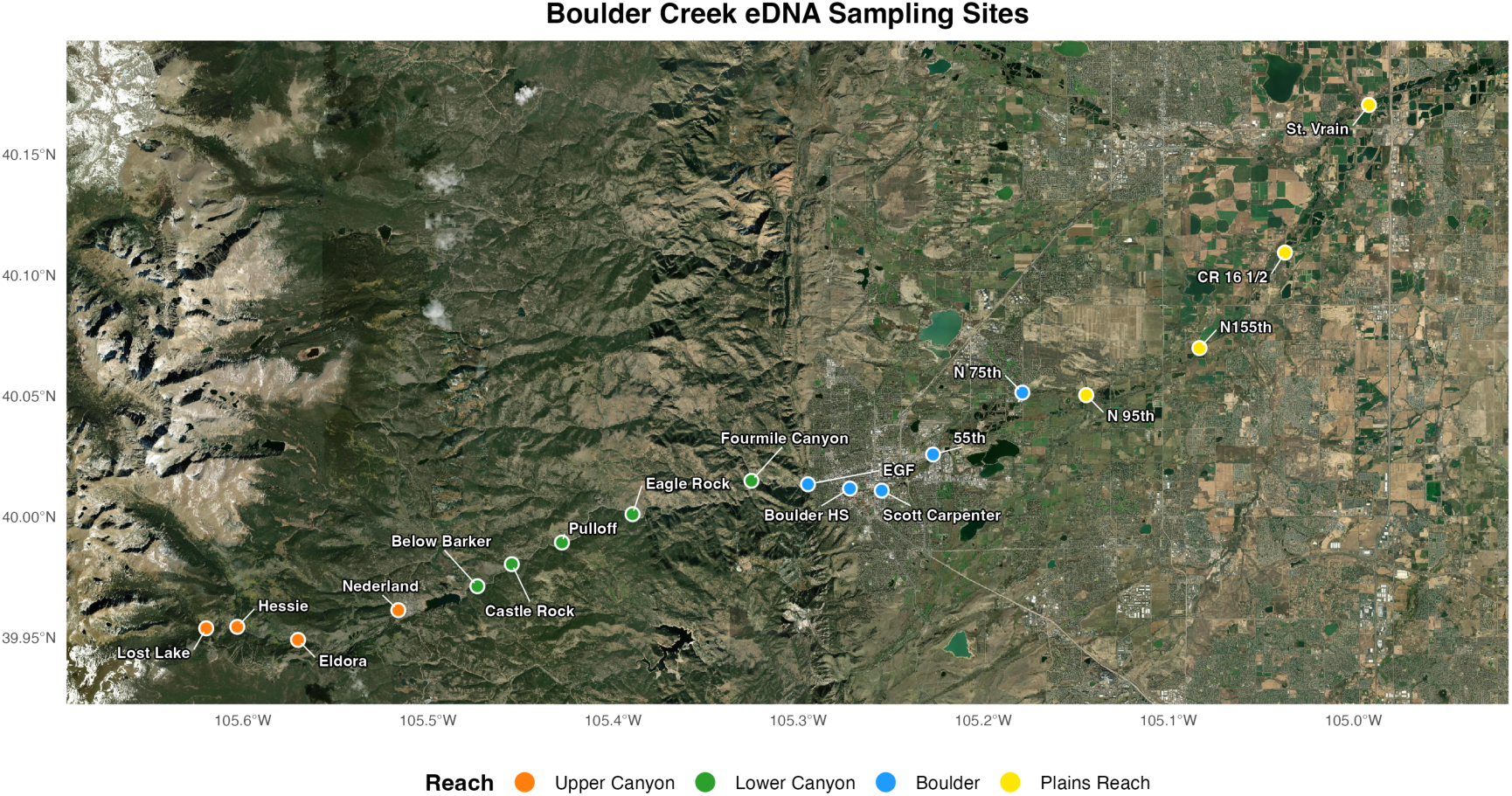
eDNA Sampling Site Map. Boulder Creek eDNA sampling sites. Points colored by reach (Upper Canyon, Lower Canyon, Boulder, Plains Reach), illustrating the longitudinal layout of the watershed from the Front Range canyon onto the plains.

### Field sampling

Three independent environmental DNA water samples were collected at each sampling site using complementary filtration approaches as part of a separate examination of filtering techniques. The first sample was collected by syringe filtration through a 25-mm diameter, 1-µm nylon membrane filter. The second sample was collected using a Jonah Ventures Baleen water collection bag, after which water was filtered through a 47-mm diameter, 1-µm nylon membrane housed in a custom 3D-printed filter housing using a portable vacuum pump. The third sample was collected by filtering water directly from the stream through an identical 47-mm, 1-µm nylon membrane mounted at the end of a telescoping sampling pole using a custom filter holder and portable vacuum pump. One site (below Barker Dam) had four samples, two of which were collected with the same gear (47-mm filter on a pole); the lower-richness member of this duplicate-gear pair was removed so that every site contributed exactly three samples, one per collection technique. For all three methods, filtration continued until flow declined below approximately 10 mL min⁻¹ due to filter clogging. Filtered volumes were recorded for every sample. Syringe filters were preserved in Longmire’s preservation buffer, whereas 47-mm filters were preserved in molecular-grade ethanol until DNA extraction. Filters were stored at ambient temperature until returning to the lab later that day and then stored frozen at –20°C until processed.

### DNA extraction

Environmental DNA was extracted from all filters using the Omega Biotek Mag-Bind® Blood & Tissue DNA HDQ 96 Kit (4×96 Preps) (Cat. No. / ID: M6399-01) according to the manufacturer’s protocol with minor modifications. Syringe filters preserved in Longmire’s preservation buffer were incubated at 56°C for one hour. Lysis buffer TL and Proteinase K were pushed through the filter housing, and all lysate was collected into a labeled 2mL tube. Ethanol used to preserve 47mm filters was removed and was disposed. The 47-mm filters were removed from the housing and placed in 5mL tubes containing lysis buffer TL and Proteinase K. All tubes containing lysate were placed in an incubator overnight at 56°C. Following overnight incubation, lysate was extracted using an automated protocol and completed using a Hamilton Microlab STARlet®. Multiple extraction blanks were included as controls to ensure extraction quality and cleanliness. Genomic DNA was eluted into 100µl and frozen at –20°C.

### Bait sets

Target-specific DNA oligos were designed using a custom bioinformatic pipeline developed by Jonah Ventures as described below. Target species lists were first compiled from publicly available biodiversity databases and regional occurrence records. The aquatic macroinvertebrate panel was constructed from Colorado Department of Public Health and Environment (CDPHE) benthic monitoring records together with GBIF occurrence data for aquatic arthropods and mollusks within Colorado (Division 2025). Reference COI-5P (Folmer) sequences were obtained from the Barcode of Life Data System (BOLD; 8 May 2026 release) and filtered to target species or, when unavailable, closely related congeners. Low-quality, duplicate, ambiguous, and short (<550 bp) sequences were removed, and a single representative sequence from each Barcode Index Number (BIN) was retained to maximize taxonomic coverage while minimizing redundancy. Candidate 80-bp bait sequences were generated by tiling across each representative COI sequence and screened to remove low-complexity regions, homopolymers, ambiguous bases, and common cloning motifs. Candidate baits were further evaluated for GC content, melting temperature, and potential off-target hybridization against a comprehensive database of human COI sequences in order to reduce likelihood of loss of hybridization potential due to contamination with human DNA. Baits were clustered at 95% sequence identity to reduce redundancy and were iteratively selected to achieve uniform coverage across target sequences while maximizing taxonomic representation and minimizing redundancy. Final bait sets were constrained to predefined panel sizes while preserving coverage of low-representation taxa and were appended with synthesis adapters for oligonucleotide production.

The final aquatic macroinvertebrate bait panel targeted the COI-5P barcode region of freshwater arthropods and mollusks expected to occur within the Boulder Creek watershed. Following quality filtering, the panel was constructed from 20,642 reference COI sequences representing 2,354 BINs, 1,409 species, and 853 genera, and comprised 12,000 unique 80-bp RNA baits with a mean of 9.0 baits per reference sequence and an average tiling density of 1.08×. Bait GC content and melting temperatures were optimized to promote uniform hybridization performance, while all baits were required to exhibit ≤84% sequence similarity to human COI to minimize off-target capture. Mean GC content was 36.1%, mean melting temperature was 66.3°C, and mean similarity to human COI was 71.0% for the macroinvertebrate panel. 80-mer targets were modified to allow for amplification and transcription and oligo pools were ordered from Twist Bioscience (South San Francisco, CA).

### Baits pool preparation

DNA oligos were amplified using Q5^®^ High-Fidelity 2X Master Mix (New England Biolabs [NEB], Ipswich, MA), purified with Monarch^®^ Spin PCR & DNA Cleanup Kit (NEB) and excess 3’ sequence was trimmed using EcoRI-HF (NEB). Trimmed oligos were transcribed using HiScribe® T7 Quick High Yield RNA Synthesis Kit (NEB) with the addition of biotinylated UTP (ThermoFisher Scientific, Waltham, MA). RNA products were cleaned using Monarch® Spin RNA Cleanup Kit (NEB) and diluted to appropriate concentrations.

### Library preparation, targeted capture, and sequencing

Genomic DNA extracted from each environmental DNA sample was converted into dual-indexed sequencing libraries using the NEBNext Ultra II DNA Library Prep Kit without DNA fragmentation to preserve long environmental DNA fragments for Oxford Nanopore sequencing. Libraries were end repaired, ligated to unique dual-index adapters, bead purified, and PCR amplified using Q5 High-Fidelity polymerase. Libraries were PCR-amplified for 18 cycles prior to hybridization.

Hybridization reactions were performed for the macroinvertebrate bait panel using the custom biotinylated RNA bait sets synthesized in-house from the bait designs described above.

Macroinvertebrate libraries were enriched using the Boulder Creek invertebrate bait panel (Pool 6.1). Each enrichment reaction used 275 ng of RNA baits and was hybridized at 65°C for approximately 16 h. Following hybridization, bait-target complexes were recovered using streptavidin-coated magnetic beads, washed under high-stringency conditions, and enriched libraries were amplified with 20 PCR cycles utilizing adapter sequences as priming sites prior to sequencing. Sequencing pool was prepared with Ligation Sequencing Kits (SQK-LSK114) according to manufacturer’s instructions and then sequenced on an Oxford Nanopore Technologies PromethION 2 Solo to a depth of 60M reads passing filter.

### Bioinformatic processing

Sequence data from the sequencing run were assembled *de-novo* into COI contigs using a custom targeted capture bioinformatics workflow. Taxonomic assignments were generated using both full assembled COI contigs compared to a full-length COI reference database, and the Folmer barcode region extracted from each contig, which was compared to a Folmer-region reference set. For invertebrate analyses, assembled contigs were retained only when the Folmer-region sequence exhibited at least 85% identity to a reference sequence. Most analyses were restricted to Arthropoda, Annelida, Mollusca.

Genus-level assignments were resolved hierarchically. The Folmer (COI) hit was preferred as the source of the genus name. Where the Folmer hit resolved only to a higher rank, the genus was taken from the standard barcode assignment. Because a contig could pass the 85% Folmer identity threshold yet inherit its genus from a lower-confidence standard-barcode hit, the source actually supplying the genus was required to meet its own quality criterion: a genus was accepted only if (i) the percent identity of the genus-supplying hit was at least 85%, or (ii) the genus came from a standard-barcode hit below 85% identity but its standard-barcode family matched the Folmer family (a family-agreement rescue). This rule removed cross-source misassignments— for example, a king-crab genus (*Paralithodes*, Arthropoda) drawn from a 73% standard-barcode hit and attached to a contig whose only confident (88% Folmer) assignment reached Gastropoda—while retaining well-supported standard-barcode genera (e.g., *Asellus*, *Choroterpes*) whose family was concordant between the two regions. Contigs whose genus failed these criteria were not discarded: the unsupported genus was set to missing and the contig was retained at its highest confidently assigned rank (its Folmer order and family at 85% identity or greater). Such contigs therefore contributed to family– and order-level richness but were excluded from genus-level analyses where a genus is required.

### Diversity metrics

Sample-level contig richness was calculated as the number of unique contigs detected within each sample. Mean richness for each sampling site was calculated by averaging replicate samples at a site. Total invertebrate richness was calculated as richness across the constituent three phyla for the union of the three samples at a site. As an indicator of biological condition, the EPT/(EPT+Chironomidae) richness proportion was calculated as the richness of Ephemeroptera, Plecoptera, and Trichoptera (EPT) contigs divided by the combined richness of EPT and Chironomidae contigs, yielding a value bounded between 0 and 1. Higher values of this index indicate greater dominance of low-water-quality-sensitive EPT taxa.

To further evaluate longitudinal shifts in assemblage sensitivity, we compiled a single master list of taxa that decrease in relative abundance with declining water quality (i.e. “decreasers”) and those that increase in relative abundance with declining water quality (i.e. “increasers”) from the State of Colorado’s macroinvertebrate multimetric index, combining the taxa designated for all three biotype regions (mountains, transition, and plains) into one composite list rather than applying region-specific designations (Colorado Department of Public Health and Environment (2017)). After combining across regions and removing duplicates, the list comprised 14 increaser taxa (families and genera including Chironomid genera such as *Cricotopus* and *Orthocladius*, and pollution-tolerant families such as Naididae and Physidae) and 30 decreaser taxa (predominantly EPT families and sensitive Chironomid and Diptera genera). Lebertiidae (an aquatic mite considered to be an increaser) was excluded from the increaser list because a single large outlier detection at one site dominated the read totals and obscured the broader gradient.

For each contig, we matched its taxonomy to the master list at genus rank where a listed genus was assigned and at family rank otherwise, and summed sequence reads for all matching contigs within each increaser or decreaser group per site. To avoid double-counting when both a family and one of its constituent genera appeared on the list, each contig was counted only once per group. We then quantified longitudinal trends in the summed read abundance of each group against river distance from Lost Lake using Spearman rank correlations and log₁₀-linear regressions, and summarised the balance between the two groups as the decreaser proportion [decreasers / (increasers + decreasers)], which we regressed against river distance.

### Sampling completeness

To assess how completely the three replicate samples per site captured invertebrate genus diversity, sample-based genus accumulation curves were constructed for each site, restricted to the three target phyla (Annelida, Arthropoda, and Mollusca). For each site, contigs were aggregated to genus level and each replicate sample was reduced to the set of genera it detected. The cumulative number of unique genera was then computed for one, two, and three pooled samples, exactly averaged over all six orderings of a site’s three samples to remove dependence on sample order. Per-step accumulation values were then averaged across the 18 sites, and the mean number of new genera added by each successive sample, together with the percentage of the three-sample genus total recovered, was used to evaluate whether three samples adequately characterized site-level genus richness.

### Spatial analyses

River distance was approximated by cumulatively summing Haversine distances between adjacent sampling locations, with Lost Lake designated as the origin. Elevation values were obtained from Google Earth. Relationships between diversity metrics and longitudinal position along the river were evaluated using an information-theoretic model selection framework.

Linear, quadratic, continuous segmented, and discontinuous segmented regression models were fit for each response variable and compared using the small-sample corrected Akaike Information Criterion (AICc). For segmented models, breakpoints were initially estimated using the *segmented* package and subsequently refit as independent linear regressions above and below each breakpoint, allowing both slope and intercept to vary between segments. This discontinuous formulation permits abrupt ecological transitions rather than requiring continuous responses along the watershed. Although the best-supported model (lowest AICc) is reported for each response (Table S2), discontinuous segmented regressions are shown consistently in figures to facilitate comparison among taxonomic groups, including mollusks for which the linear model was best supported.

### Assemblage analyses

Assemblage structure was visualized using non-metric multidimensional scaling (NMDS) based on Jaccard dissimilarities for all contigs. Associations between NMDS scores and river distance were evaluated using Spearman rank correlations, while differences among watershed reaches were tested using permutational multivariate analysis of variance (PERMANOVA; 999 permutations). Similarity percentage (SIMPER) analysis was used to identify genera contributing most strongly to compositional differences among reaches. Assemblage turnover along the watershed was quantified using pairwise Jaccard similarity as a function of river distance.

Distance-decay relationships were evaluated using log-linear regression and Mantel tests (999 permutations). An additional nested-model F-test evaluated whether assemblage similarity declined more rapidly across the canyon-to-plains transition than expected from geographic distance alone.

### Sequence characteristics

Mean assembled contig length and mean read abundance per contig were calculated for each site using Arthropoda, Annelida, and Mollusca detections. Site-level means were regressed against river distance, and associations were quantified using Spearman rank correlations. Contig-length distributions were summarized separately for assembled (untrimmed) sequences and Folmer-trimmed barcode regions.

### Comparison with conventional bioassessment

Morphological macroinvertebrate data were obtained from the Colorado Department of Public Health and Environment Multimetric Index (MMI) monitoring program (Colorado Department of Public Health and Environment 2024). Arthropod richness was calculated after excluding non-arthropod taxa while retaining aquatic mites, crustaceans, and collembolans to match the arthropod definition used for eDNA analyses. Data from Boulder Creek and Middle Boulder Creek sites were included to match eDNA sampling.

Longitudinal richness patterns for eDNA and MMI datasets were compared using separate linear regressions for canyon and Boulder–Plains reaches. Average arthropod richness among sites was additionally summarized by reach (canyon sites and sites below the canyon) for direct comparison between methods. To ensure comparable spatial coverage, canyon eDNA summaries were restricted to sites either between the two MMI canyon sampling locations or adjacent to them.

### Shotgun metagenomic comparison

To determine enrichment levels from hybridization, we sequenced the original metagenomic pools to similar sequencing depths as the targeted capture pools and then determined the percentage of all reads that hit our target species using the same techniques for taxonomic profiling as targeted capture pools. Enrichment magnitude was calculated as the fold-change between paired un-enriched shotgun samples and enriched targeted capture samples, in the proportion of raw reads for a given sample that contained arthropod COI. For a given sample enrichment was calculated as the ratio of the percentage of target COI in the enriched sample to the unenriched sample.

### Statistical analyses

All statistical analyses were performed in R version 4.4.2 (R Core Team 2024). Unless otherwise indicated, statistical significance was assessed at α = 0.05.

Analyses and figures were produced using the **tidyverse** suite (Wickham et al. 2019), **segmented** (Muggeo 2008), **vegan** (Oksanen et al. 2025), **patchwork**, and **gt**. Package versions: **vegan** 2.6.10, **segmented** 2.1.4, **ggplot2** 4.0.2, **patchwork** 1.3.0, **dplyr** 1.2.0, **tidyr** 1.3.1, **readr** 2.1.5, **stringr** 1.5.1, and **gt** 1.0.0.

## Results

### Sequencing performance and taxonomic assignment

Targeted capture enriched COI sequences for the three phyla (Arthropoda, Annelida, and Mollusca) by a median of 1,760-fold across 54 samples. Across the Boulder Creek samples enriched with the invertebrate bait set, COI-containing reads represented a median of 1.9% of all raw reads (spanning 0.6-21% among samples), compared with a median of only 0.0011% in the paired shotgun metagenomic libraries prepared from the same eDNA. Across the 54 Boulder Creek invertebrate targeted-capture libraries, sequencing yielded a median of 174,490 demultiplexed reads per sample (range 20,717–532,066). A median of 15,237 reads mapped to the reference database prior to filtering (7.9% of demultiplexed reads), and 8,182 mapped reads remained after further quality filtering (4.7% of demultiplexed reads). These filtered mapped reads formed the basis for downstream contig assembly and taxonomic assignment. Across all samples, contig assembly yielded 2,678 contig sequences assigned to Arthropoda, Annelida, or Mollusca, of which 2,301 (85.9%) met the criteria for confident genus-level assignment; the remaining contigs were retained at the family or order level.

Targeted capture recovered relatively long COI contigs (**Figure 2**). Targeted capture detected an average of 49 ± 32 (s.d.) invertebrate contigs per sample. Assembled untrimmed contigs averaged 452 bp, with nearly one-quarter of all contigs (24.6%) exceeding 658 bp, the typical length of the Folmer COI barcode. Following trimming to restrict contigs to the Folmer region, macroinvertebrate contigs averaged 406 bp (median = 348 bp), with 64.5% at least 300 bp in length. Mean contig length increased with increasing distance from Lost Lake (Spearman ρ = 0.62, *p* = 0.007), rising by 1.2 bp per km (linear regression: *R*² = 0.36, *p* = 0.009)—an increase of roughly 76 bp across the ∼63 km gradient—whereas read depth per contig was invariant with distance (**Figure S1**).

**Figure 2.**
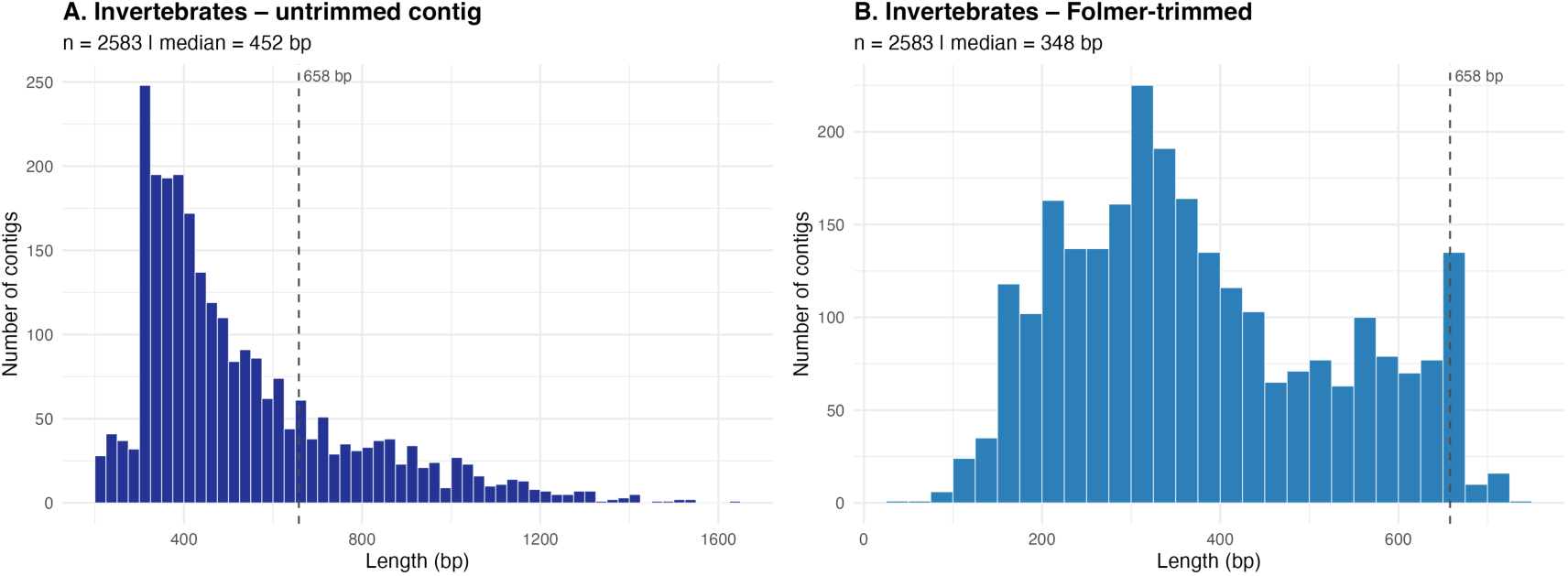
Sequence Length Distributions: Untrimmed vs Folmer-trimmed Contigs. Univariate length distributions for invertebrate contigs (three main phyla: Arthropoda, Annelida, Mollusca), shown for untrimmed assembled contigs (left column) and Folmer-trimmed barcodes (right column). Dashed line marks the Folmer COI barcode length (∼658 bp).

Recovered sequences exhibited high similarity to reference databases within the Folmer COI barcode region. Among the fifteen most frequently detected macroinvertebrate genera, sequence identity averaged 97.8%, with individual genera ranging from 88.2% (*Sulcospira*) to 99.7% (*Hydropsyche*). Most dominant aquatic insect genera—including *Baetis*, *Hydropsyche*, *Ephemerella*, and *Sweltsa*—were represented by contigs exceeding 400 bp and sequence identities greater than 98.9%, supporting confident taxonomic assignment (**Table 1**).

**Table 1.** Most Frequently Observed Invertebrate Genera. The 15 most frequently detected invertebrate genera across samples, with Folmer COI region percent-identity statistics and mean Folmer-trimmed sequence length. ‘Samples’ and ‘Sites’ are the number of distinct samples and sites a genus was detected in; ‘Mean %ID’ is the mean percent identity of the Folmer-region match; ‘Folmer len (bp)’ is the mean length of the Folmer-trimmed sequence.

| Phylum | Genus | Samples | Sites | Contigs | Mean %ID | %ID range | Folmer len (bp) |
| --- | --- | --- | --- | --- | --- | --- | --- |
| Arthropoda | *Baetis* | 48 | 18 | 90 | 99.3 | 90.7–100.0 | 474 |
| Arthropoda | *Micropsectra* | 36 | 17 | 78 | 97.7 | 87.9–100.0 | 373 |
| Arthropoda | *Simulium* | 36 | 16 | 122 | 98.2 | 85.0–100.0 | 357 |
| Arthropoda | *Cricotopus* | 32 | 13 | 119 | 97.4 | 89.1–100.0 | 394 |
| Arthropoda | *Ephemerella* | 31 | 12 | 64 | 98.9 | 86.5–100.0 | 454 |
| Arthropoda | *Orthocladius* | 31 | 14 | 59 | 98.2 | 90.9–100.0 | 353 |
| Arthropoda | *Hydropsyche* | 30 | 14 | 46 | 99.7 | 98.0–100.0 | 404 |
| Arthropoda | *Sweltsa* | 28 | 12 | 37 | 99.5 | 94.8–100.0 | 428 |
| Annelida | *Nais* | 27 | 12 | 93 | 98.3 | 86.5–100.0 | 387 |
| Annelida | *Chaetogaster* | 24 | 13 | 61 | 98.4 | 87.9–100.0 | 436 |
| Arthropoda | *Eukiefferiella* | 24 | 13 | 51 | 96.4 | 89.6–100.0 | 408 |
| Mollusca | *Sulcospira* | 23 | 13 | 36 | 88.2 | 85.0–91.4 | 282 |
| Arthropoda | *Daphnia* | 21 | 8 | 46 | 99.7 | 98.0–100.0 | 382 |
| Arthropoda | *Bosmina* | 20 | 8 | 37 | 99.6 | 94.2–100.0 | 440 |
| Arthropoda | *Tricorythodes* | 20 | 8 | 31 | 99.2 | 94.6–100.0 | 391 |

### Detected taxa

Targeted capture recovered a diverse invertebrate assemblage spanning 388 genera, 191 families, and 43 orders across the three target freshwater phyla (Arthropoda, Annelida, and Mollusca).

Incorporating detections from five additional non-target invertebrate phyla (Porifera, Cnidaria, Platyhelminthes, Nematoda, and Bryozoa) raised the total to 460 genera across eight phyla (**Figure 3**). Arthropods accounted for the majority of the recovered diversity, comprising 345 genera distributed among 171 families and 33 orders, while Annelida contributed 30 genera from 9 families and 6 orders, and Mollusca contributed 13 genera representing 11 families and 4 orders. Within Arthropoda, Diptera was the most diverse order (128 genera), driven largely by the non-biting midges (Chironomidae, 67 genera). EPT orders together contributed 53 genera (Ephemeroptera, 18; Plecoptera, 17; Trichoptera, 18) while Coleoptera added a further 40 genera. Beyond the aquatic insect fauna, targeted capture recovered substantial diversity of other riparian and terrestrial arthropods, including 27 genera of Lepidoptera, most of which were terrestrial but also included the aquatic moth genus *Petrophila*, 21 arachnid genera, predominantly spiders (Araneae), and 8 genera of springtails (Collembola), reflecting the deposition of terrestrial biodiversity into stream eDNA. Among the non-arthropod taxa, annelid diversity (30 genera) included both aquatic oligochaetes (e.g. *Nais*, *Chaetogaster*, and *Tubifex*) and terrestrial oligochaetes (e.g. *Lumbricus* and *Enchytraeus*), while molluscan detections (13 genera) included freshwater snails and bivalves such as *Physella*, *Corbicula*, and *Pisidium*.

**Figure 3.**
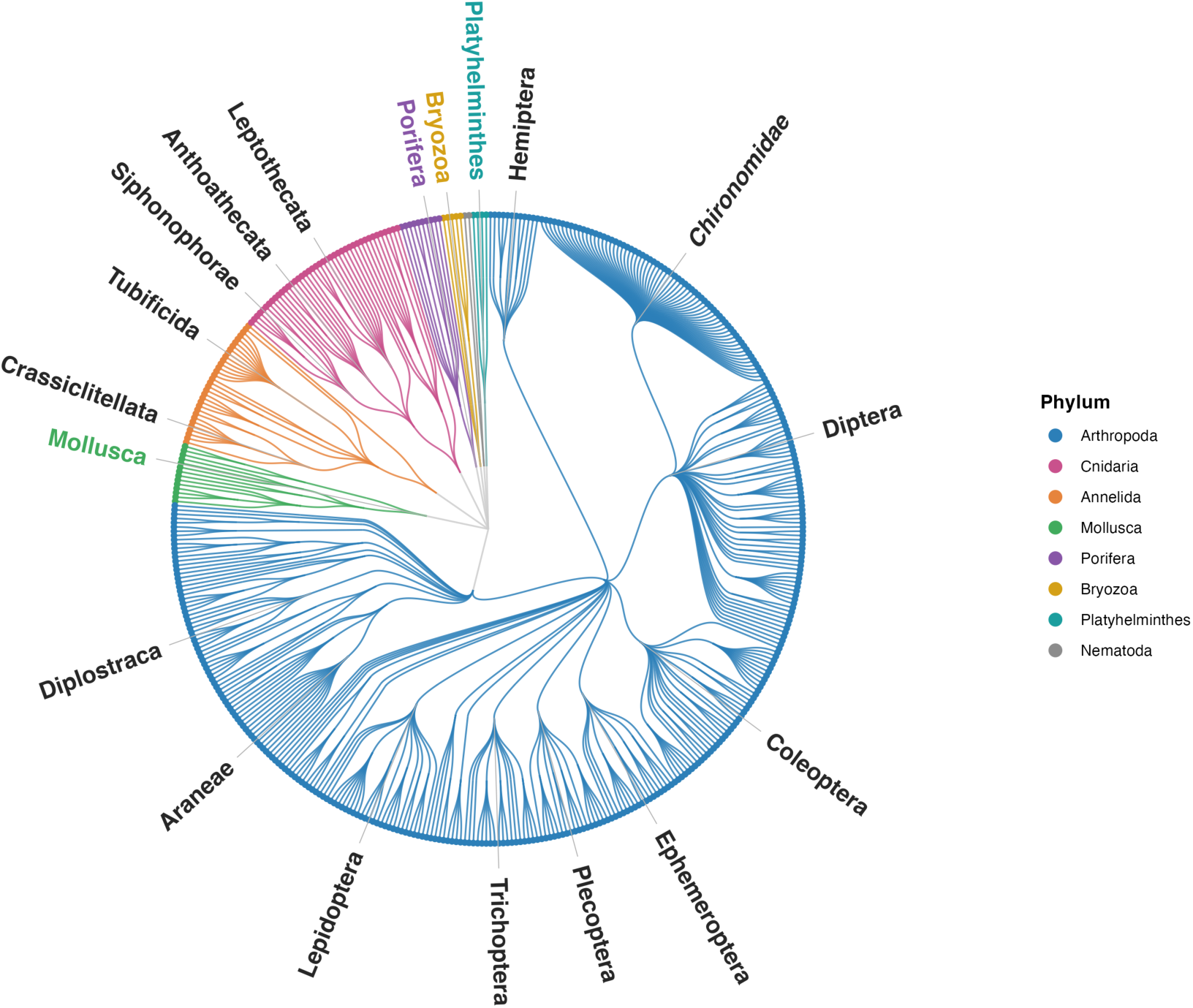
Invertebrate Taxonomic Tree. Circular dendrogram of the eDNA-detected invertebrate fauna, arranged by taxonomy (phylum → class → order → family → genus). Each outer tip is a genus (460 genera across 8 phyla); branches and tips are colored by phylum. Labels mark the major orders (≥ 8 genera) plus the family Chironomidae; all other internal ranks are left unlabeled for clarity. The three focal phyla (Arthropoda, Annelida, Mollusca) are drawn from the vetted genus-level community, and five additional detected phyla (Porifera, Cnidaria, Platyhelminthes, Nematoda, Bryozoa) are included from the broader detection table.

Sequences from a snail that best matched *Sulcospira*—a Southeast Asian genus—at less than 90% identity were also detected, though its true identity remains uncertain.

The most widespread detected invertebrate genera are commonly found in local streams (**Figure 4**; **Table 1**) and spanned all three phyla. Mayflies in the genus *Baetis* were detected at all 18 sampling sites and in 48 of the 54 water samples, making it the most frequently observed genus overall. Among Diptera, the chironomid genera *Micropsectra*, *Orthocladius*, and *Cricotopus* were detected at 17, 14, and 13 sites, respectively, while the black fly *Simulium* occurred at 16 sites. Other frequently detected aquatic insect genera included the mayfly *Ephemerella* (12 sites), the caddisfly *Hydropsyche* (14 sites), and the stonefly *Sweltsa* (12 sites), representing all three EPT orders. Common non-insect taxa included the annelid genera *Nais* and *Chaetogaster*, detected at 12 and 13 sites, respectively, and the freshwater snail best matching *Sulcospira*, which was recovered from 13 sites.

**Figure 4.**
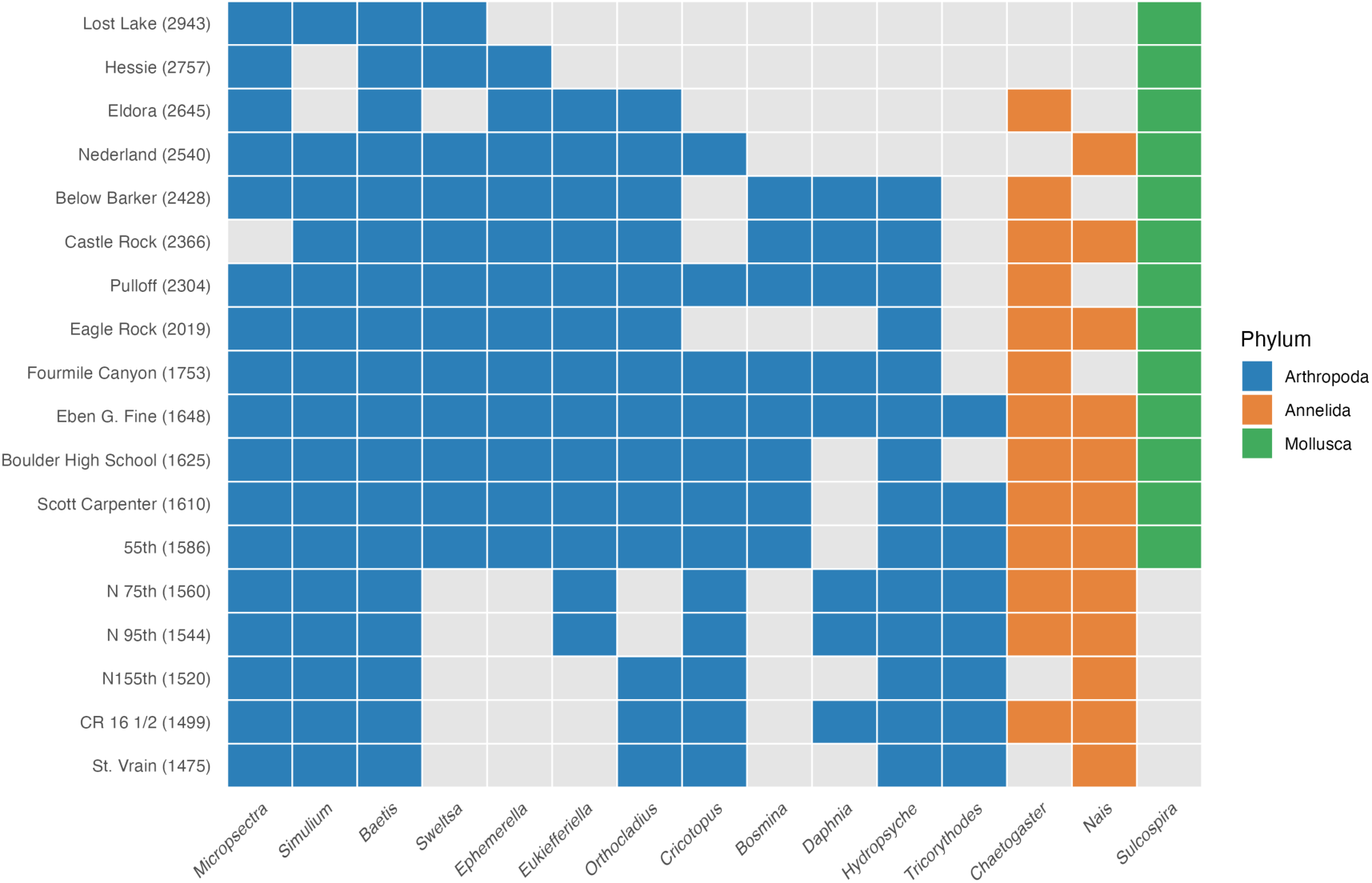
Top 15 Most Frequently Observed Invertebrate Genera. Presence/absence heatmap of the 15 most frequently detected invertebrate genera (three main phyla: Arthropoda, Annelida, Mollusca) across sites. Sites (y-axis) are ordered by elevation (m elevation, highest at top); genera (x-axis) are ordered within phylum by first appearance (the highest-elevation site at which each genus is detected). Cells are colored by phylum where the genus is present and grey where absent.

Taxa accumulation curves demonstrated substantial gains in invertebrate taxonomic recovery with replicate sampling (**Table S4**). A single water sample recovered an average of 27.5 invertebrate genera per site, representing 48.8% of the total genera detected across three replicates. A second sample contributed an additional 16.1 genera, increasing cumulative richness to 43.6 genera (77.4% of the three-sample total). The third replicate added a further 12.7 genera, yielding a mean of 56.4 genera per site and demonstrating that even after two water samples, nearly one-quarter of the detectable genera remained unsampled. Although mean filtered volumes differed substantially among collection approaches (297 ± 158 mL for the 25-mm syringe filter, 752 ± 345 mL for the 47-mm baleen sampler, and 656 ± 453 mL for the 47-mm pole sampler), no significant differences in arthropod or annelid richness were observed among these sampling approaches (Kruskal–Wallis, *P* > 0.4 for both groups) (**Table S5, Table S6**).

### Longitudinal patterns in macroinvertebrate diversity and bioassessment indices

Macroinvertebrate richness exhibited strong longitudinal structure along Boulder Creek (**Figure 5**) with distinct ecological transitions near the canyon-to-plains boundary. Total richness increased progressively through the upper and lower canyon before reaching maximum values within the Boulder reach, followed by a marked decline downstream through agricultural reaches. A discontinuous segmented regression provided the best-supported model (tied for lowest AICc) and explained 48.3% of the variation in total richness (*P* = 0.023; **Table S2**).

**Figure 5.**
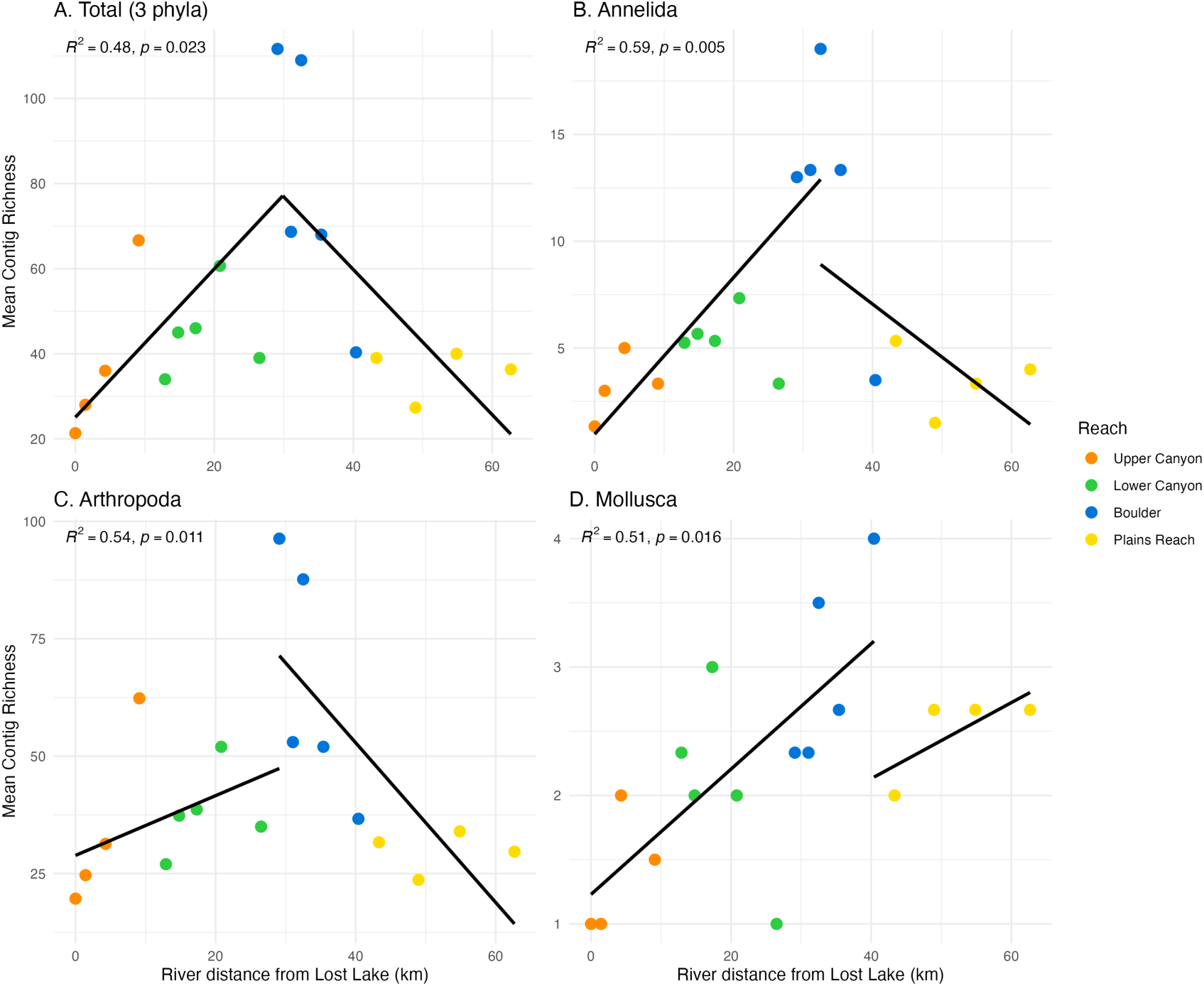
Invertebrate Contig Richness vs River Distance. Total richness (A) and Annelida (B), Arthropoda (C), Mollusca (D) richness as a function of distance from the highest-elevation site. Discontinuous piecewise regressions shown for each panel.

Similar longitudinal patterns were observed independently for Arthropoda, Annelida, and Mollusca, although each taxonomic group exhibited distinct rates of increase and downstream decline. Arthropod richness displayed the strongest spatial pattern, increasing steadily through the canyon before declining below Boulder, with the discontinuous segmented model explaining 54% of the variation (*P* = 0.011). Annelid richness increased more gradually and remained elevated across portions of the urban corridor (R² = 0.59; *P* = 0.005). Molluscan richness increased through the canyon and into Boulder, before declining in the plains reach (R² = 0.51; P = 0.016).

Among bioassessment indicator taxa, richness of EPT taxa increased through the canyon before declining sharply downstream of Boulder (**Figure 6**). A discontinuous segmented regression explained 57.2% of the variation in EPT richness (*P* = 0.007), with a breakpoint near the canyon-to-plains boundary. In contrast to the more sensitive EPT taxa, Chironomidae richness peaked further downstream and declined only in the lowest reaches (**Figure 6**). A discontinuous segmented regression (R² = 0.37) placed the breakpoint at approximately 12 km downstream of the EPT breakpoint. The EPT/(EPT+Chironomidae) richness proportion declined significantly downstream (Spearman ρ = –0.77, *P* < 0.001; linear R² = 0.59), falling below 0.5 in downstream urban and agricultural reaches where Chironomidae richness exceeded EPT richness (**Figure 6**). Screening the broader suites of “increaser” and “decreaser” taxa used in Colorado bioassessment, 20 of the 30 listed decreaser taxa and 11 of the 14 listed increaser taxa were recovered across the watershed (**Figure 7**). These two groups showed opposing longitudinal responses. Read abundance of decreaser taxa declined sharply with river distance from Lost Lake (Spearman ρ = –0.76, *P* < 0.001, **Figure 8**), whereas read abundance of increaser taxa increased downstream (Spearman ρ = 0.63, *P* = 0.005; **Figure 8**). Consequently, the proportion of decreaser reads declined significantly downstream, from near 1.0 in the headwaters to near 0 in the plains (Spearman ρ = –0.86, *P* < 0.001; **Figure 8**).

**Figure 6.**
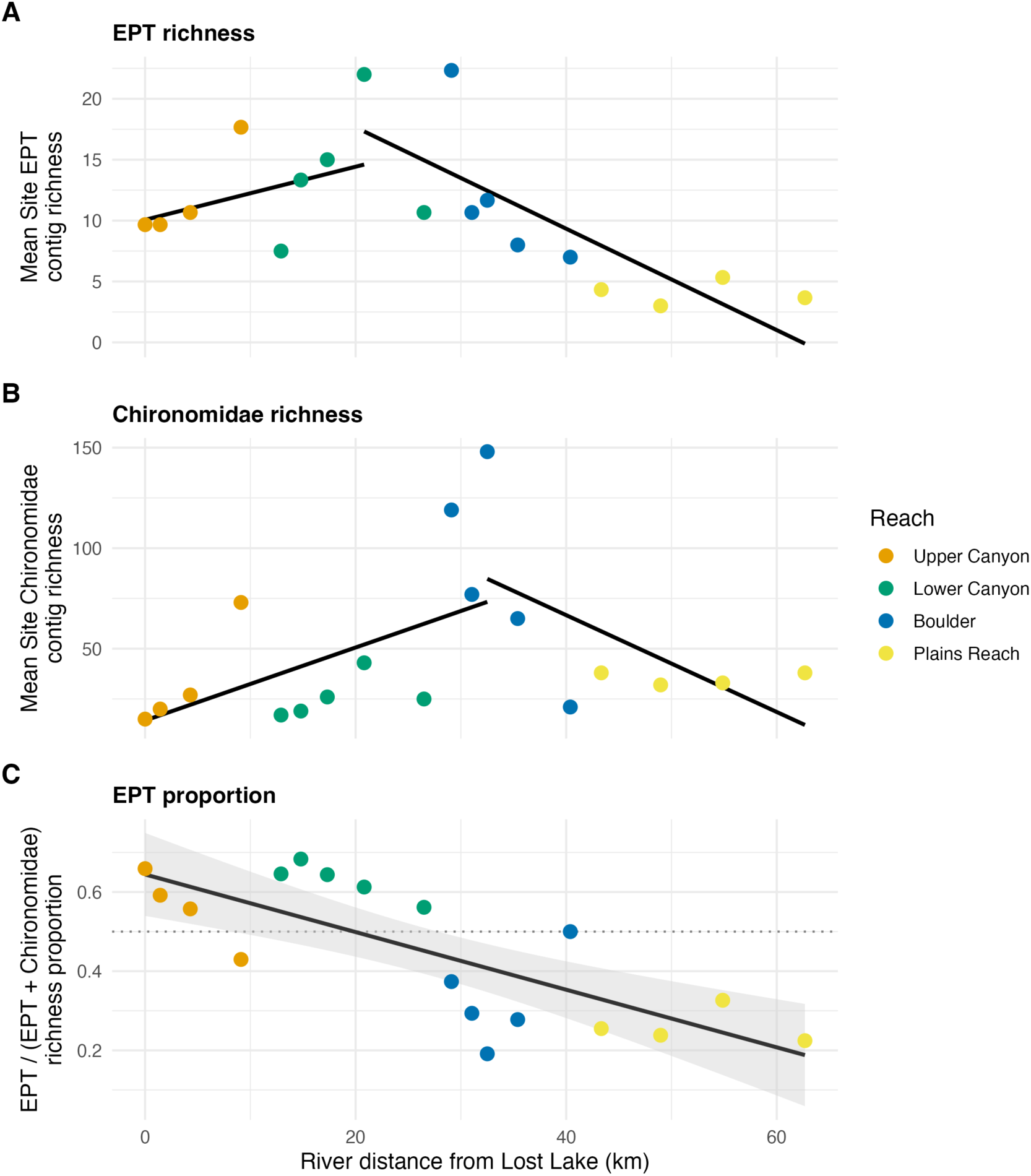
EPT Richness, Chironomidae Richness, and EPT Proportion vs River Distance. Three-panel comparison versus distance from the highest-elevation site, points colored by reach. (A) Mean EPT (Ephemeroptera, Plecoptera, Trichoptera) contig richness, with a discontinuous segmented fit (breakpoint ∼20.8 km). (B) Chironomidae contig richness, with a discontinuous segmented fit (breakpoint ∼32.5 km). (C) The EPT proportion [EPT/(EPT+Chironomidae) contig richness]. Values above the dotted line (0.5) indicate EPT dominance, shown with a linear fit (AICc-best; Table S2). Proportion: Spearman rho = –0.77, p = 0.0003; r^2^= 0.59.

**Figure 7.**
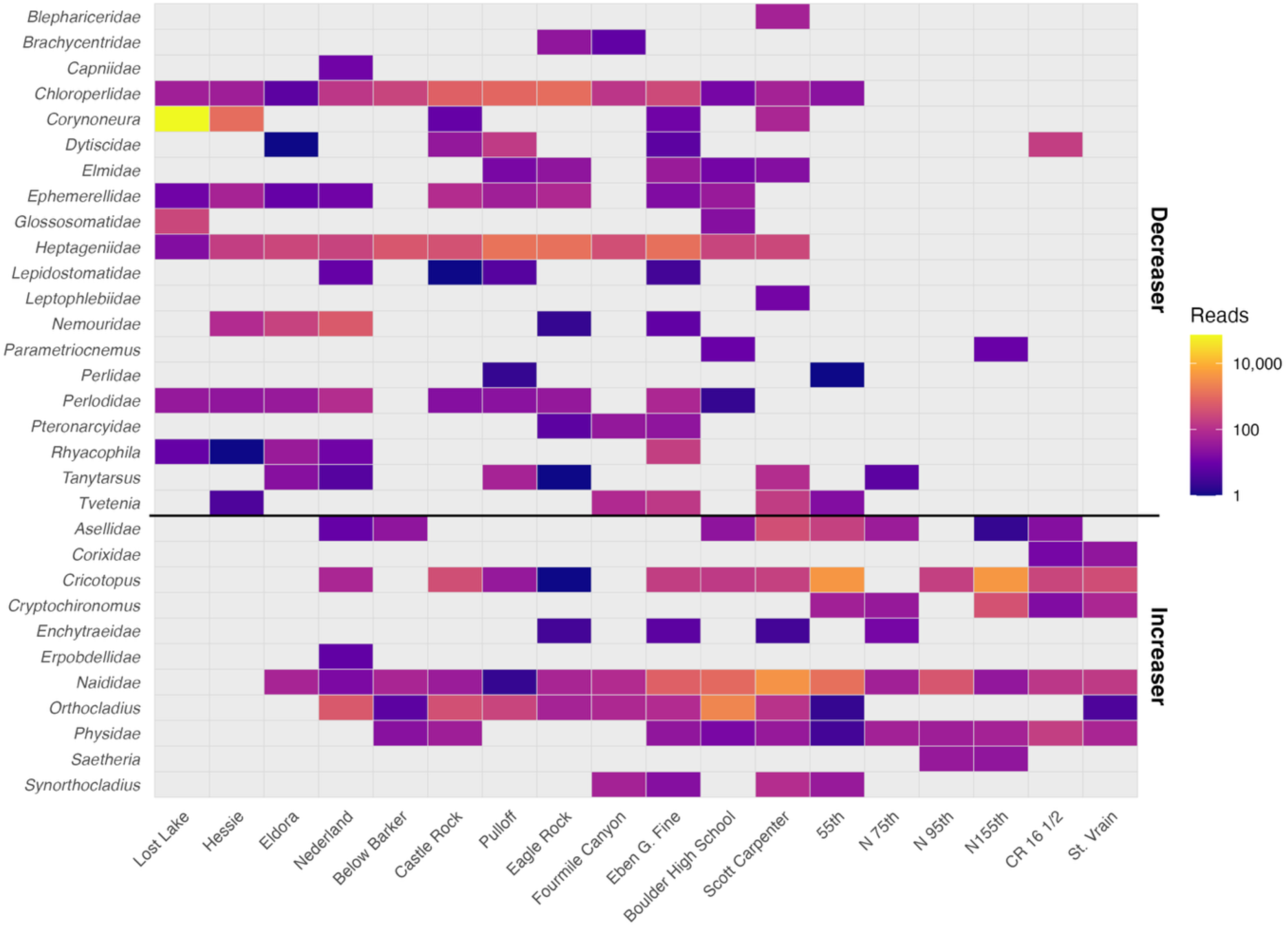
Increaser and Decreaser Taxon Read Abundance by Site. Heatmap of eDNA read abundance for Colorado bioassessment indicator taxa across the 18 Boulder Creek sites (columns, ordered by elevation). Rows are decreaser (pollution-sensitive) taxa above the black divider and increaser (pollution-tolerant) taxa below. Cells are colored by summed contig reads (grey = not detected). A contig is matched to a listed taxon by genus (preferred) or family.

**Figure 8.**
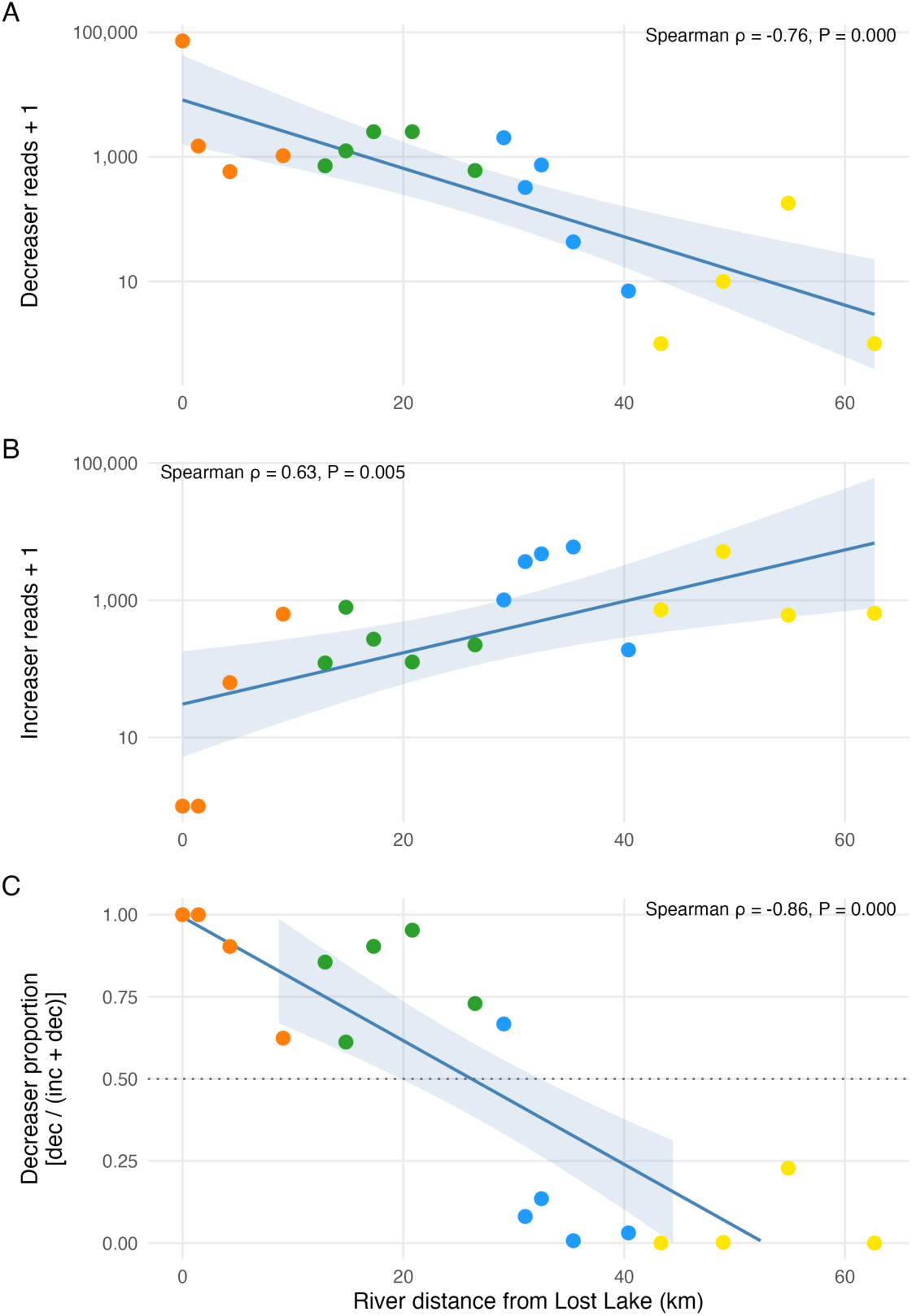
Increaser and Decreaser Reads and Decreaser Proportion vs River Distance. Longitudinal patterns in the Colorado bioassessment indicator taxa (Lebertiidae excluded) vs river distance from Lost Lake. (A) Total decreaser reads + 1 (log scale) with a log-linear fit. (B) Total increaser reads + 1 (log scale) with a log-linear fit. (C) The decreaser proportion [decreasers / (increasers + decreasers)] with a linear fit.

### Invertebrate assemblage turnover along the watershed

Macroinvertebrate assemblage composition changed progressively along the longitudinal gradient of Boulder Creek (**Figure 9**). NMDS ordination revealed a strong directional shift in assemblage composition from headwater to downstream sites, with NMDS axis scores strongly correlated with river distance (Spearman ρ = 0.90, *P* < 0.001). Reach category explained a substantial proportion of assemblage variation (PERMANOVA R² = 0.41, *P* = 0.001). Ordering sites by river distance traced a coherent downstream trajectory through ordination space, progressing from upper-to lower-canyon assemblages through the Boulder reach before a pronounced shift into the distinct plains-reach cluster, with the largest compositional shift occurring at the canyon-to-plains transition (**Figure 9**). There was no significant change in the total sum of sequence reads from contigs assigned to the three main invertebrate phyla with increasing distance from Lost Lake (Spearman ρ = 0.05, *P* = 0.84; **Figure S2**), nor was there any trend in sequencing effort (ρ = –0.09, P = 0.71). Pairwise Jaccard similarity decayed with river distance (Mantel r = 0.79, P = 0.001), halving roughly every 25 km, and site pairs spanning the canyon-to-plains transition were less similar than distance alone predicts (nested-model F-test, P = 0.002)—independently confirming the discontinuity seen in the ordination (**Figure S3**).

**Figure 9.**
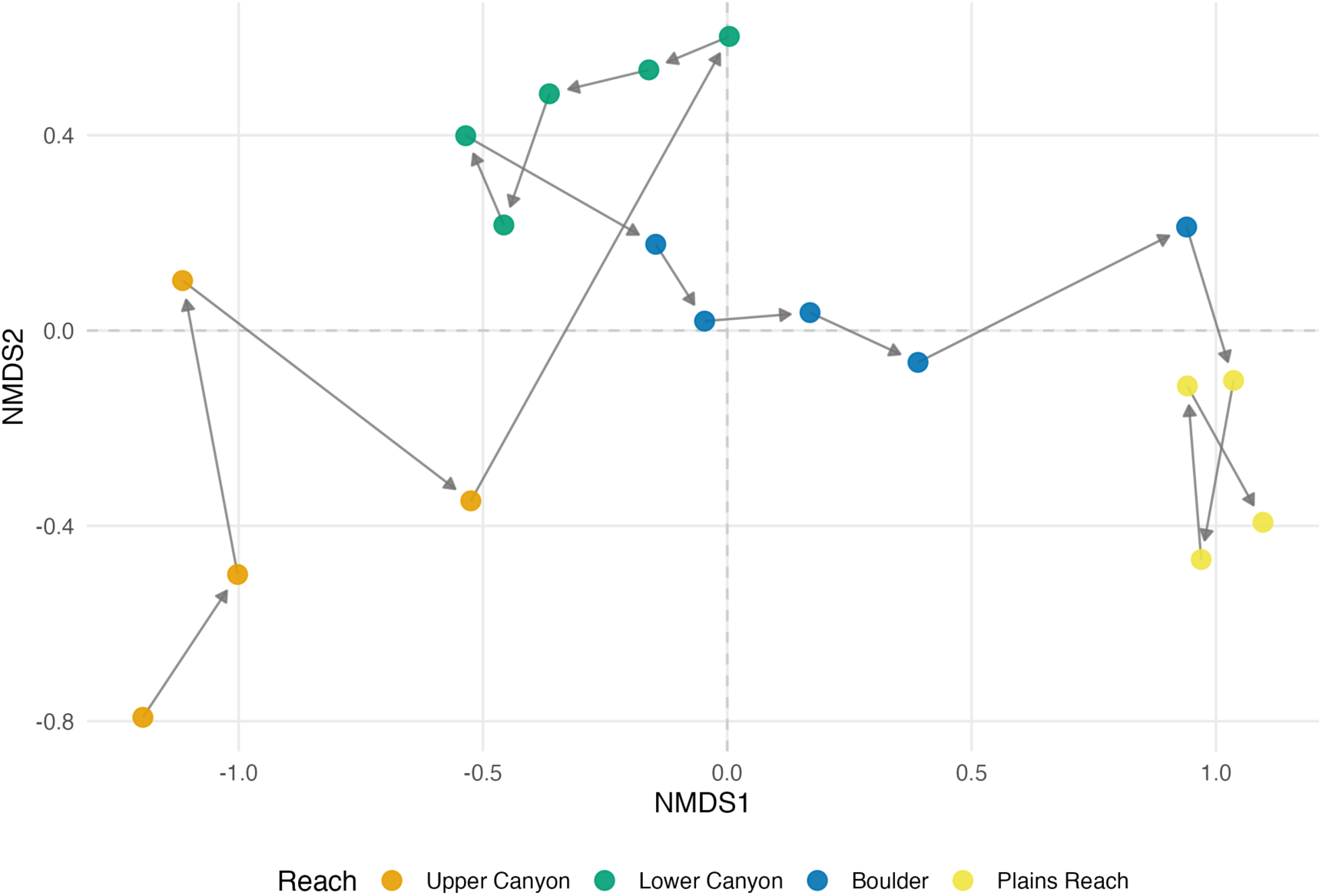
Downstream Trajectory of Invertebrate Community with NMDS Ordination. Genus-level Jaccard NMDS ordination of invertebrate communities (stress = 0.087), plotting NMDS1 against NMDS2. Points colored by reach category. Grey arrows connect sites with decreasing elevation, tracing the upstream-to-downstream succession through ordination space.

SIMPER analysis identified distinct genera contributing to community turnover among stream reaches (**Table S7**). Upper canyon communities were characterized by cold-water taxa including the mayfly *Cinygmula*, the stonefly *Paraleuctra*, the caddisfly *Rhyacophila*, and the freshwater snail currently identified as *Sulcospira*. Beginning in the lower canyon, assemblages shifted toward taxa including *Hydropsyche*, which remained common through Boulder and the plains, together with cladoceran genera such as *Bosmina* and *Daphnia*. Boulder reaches were further distinguished by increased occurrence of *Psectrocladius*, *Eukiefferiella*, *Boreoheptagyia*, and *Acanthocyclops*. Plains reaches were distinguished by genera typical of larger, lower-elevation streams, including the mayflies *Fallceon* and *Tricorythodes* and the chironomid *Dicrotendipes*.

### Comparison with conventional bioassessment

Many of the taxa with the highest number of individuals detected with kicknetting in the State of Colorado surveys were among the most detected invertebrate taxa with targeted capture (**Table S8**). For example, individuals of the genus *Baetis* were the most detected arthropod in the kicknet data (3455 individuals) and were the third-most detected arthropod for targeted capture. Individuals of the genus *Nais* were the most detected annelid with both the kicknet and targeted capture techniques. The most detected arthropod with targeted capture (*Simulium*) was the fourth-ranked arthropod in the kicknet data. In general, most of the 15 most-detected taxa with kicknetting were also among the most detected with targeted capture. Discrepancies could also in part be explained by differences caused by poor DNA reference database, e.g. *Sulcospira* is not known to be present in the Colorado and was not listed as having been detected in the MMI data. Only two mid-elevation canyon sites were sampled by the state. *Fallceon* was the 2nd most abundant arthropod by kicknet and the 36th-ranked arthropod with targeted capture, which could be influenced by the mismatch between distributions and sampling density.

Although morphological taxon counts and contig richness or read counts are not directly equivalent, longitudinal patterns generated by targeted capture closely paralleled those obtained from the Colorado Multimetric Index (MMI) macroinvertebrate monitoring program (**Figure 10**). Despite fundamentally different sampling strategies and taxonomic workflows, both approaches showed increasing arthropod richness through the canyon, peak richness near the canyon mouth, and declining richness downstream across the Boulder and plains reaches. Within the canyon reaches where both methods overlapped spatially (**Table S9**), average arthropod richness was similar between methods. Morphological surveys averaged 42.5 taxa (range 39–46), while targeted capture averaged 41.3 arthropod contigs per sample (range 27.0–62.3). In the Boulder city and plains reaches, targeted capture recovered greater average arthropod richness (49.4) than morphology (31.9), although the richness distributions overlapped substantially (23.7–96.3 versus 18–51), indicating that both methods identified the same broad longitudinal richness pattern while targeted capture frequently recovered greater arthropod diversity.

**Figure 10.**
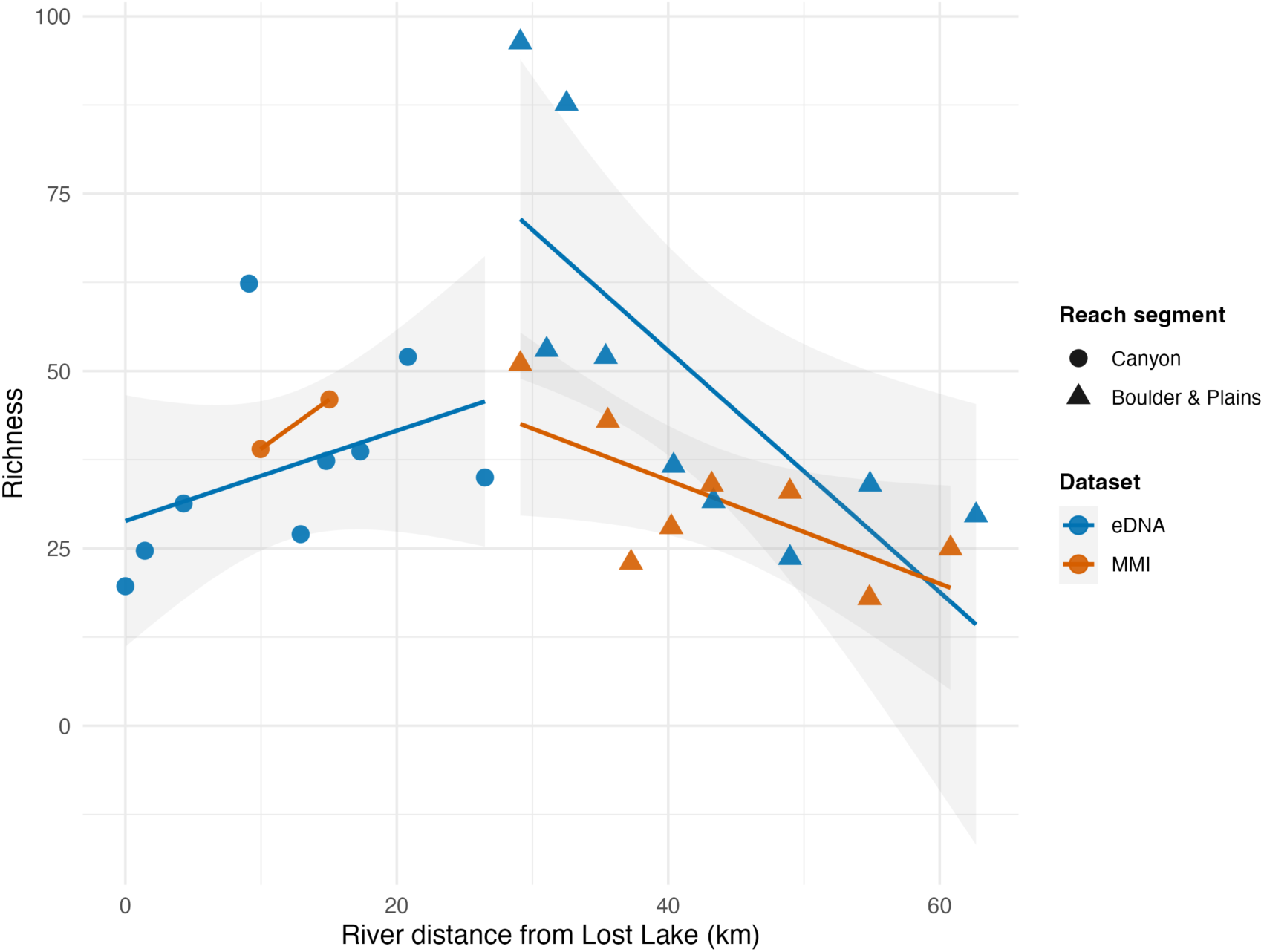
eDNA Arthropod vs MMI Arthropod Richness vs Distance from Lost Lake. eDNA arthropod contig richness (blue) and CDPHE MMI morphology-based arthropod taxa richness (orange; insects plus water mites, crustaceans, and Collembola) plotted against straight-line distance from Lost Lake. Two linear segments are fit per dataset: Canyon (circles) vs Boulder & Plains (triangles). eDNA Canyon slope = 0.64, Boulder & Plains slope = –1.70 taxa/km; MMI Canyon slope = 1.38, Boulder & Plains slope = –0.73 taxa/km.

## Discussion

### Targeted capture detects diverse assemblages of freshwater bioassessment species

This study demonstrates that targeted capture can recover ecologically informative macroinvertebrate assemblages across an entire watershed from a single COI-based workflow. Across 18 sites spanning nearly 1,500 m of elevation and multiple ecological zones, capture consistently recovered long mitochondrial contigs suitable for genus– and species-level assignment. The resulting assemblage data reproduced known longitudinal patterns—rising macroinvertebrate richness through the mountain canyon and its decline across the plains, and strong correspondence with independently developed bioassessment metrics.

A notable finding was the breadth of invertebrate diversity recovered from a relatively modest sampling effort. Nearly 400 genera across the three targeted phyla were detected watershed-wide, including groups rarely resolved comprehensively in conventional survey. Among those were 67 chironomid genera, a level of resolution difficult to achieve by morphology as larval chironomids are notoriously difficult to identify (Merritt, Cummins, and Berg 2019). It should be noted that not all of the recovered genera from the three target invertebrate phyla are aquatic. A portion, including many Lepidoptera, arachnids, and beetles, represent terrestrial or riparian taxa deposited into the stream rather than potential aquatic bioassessment species. Capture also recovered COI from several metazoan phyla absent from the bait design (Cnidaria, Bryozoa, Porifera), reflecting hybridization’s tolerance of sequence divergence, which lets probes designed from one lineage enrich homologous COI from related taxa; this breadth is not merely nonspecific enrichment, as candidate probes exceeding 84% identity to human COI were excluded. Because a single panel of many probes removes the need for universally conserved primer-binding sites, capture makes it practical to recover the COI barcode across broad freshwater assemblages in one assay—and because COI has the most comprehensive reference database and highest resolution for metazoans (Ratnasingham and Hebert 2007), it offers a framework for standardizing cross-phylum bioassessment around a single, well-characterized marker.

### Targeted capture as a distinct strategy for environmental DNA

Targeted capture represents a distinct molecular strategy for environmental DNA analysis that removes the constraints of qPCR and metabarcoding by replacing enrichment by PCR amplification with hybridization-based enrichment. Rather than requiring conserved primer binding sites, capture probes can tile across sequence diversity at one or multiple loci, allowing highly informative markers such as COI to be recovered from phylogenetically diverse freshwater communities independent of whether conserved primer sites would exist for a desired target group. Previous studies have established the feasibility of targeted capture for environmental DNA, but most have focused on technical proof-of-concept, small taxonomic inventories, or recovery of complete mitochondrial genomes. The first freshwater application was presented by Wilcox et al. (2018), who used a relatively small panel of biotinylated DNA baits targeting mitochondrial DNA from multiple freshwater phyla including Arthropoda, Chordata, Mollusca, Annelida, and Cnidaria. That effort demonstrated that hybridization capture could enrich a small set of target sequences recovered from freshwater environmental DNA rather than evaluating targeted capture as a framework for ecological assessment. Additional studies, including the vertebrate surveys of tropical waterholes (Seeber et al. 2019; Li et al. 2023) and whale shark monitoring from seawater (Jensen et al. 2021), have similarly demonstrated that target enrichment can increase recovery of informative mitochondrial sequences from complex environmental samples. Outside aquatic environmental DNA, closely related hybridization-capture approaches have become widely adopted in ancient sediment DNA research, where mitochondrial and nuclear DNA recovered from cave sediments have been used to reconstruct prehistoric human populations and associated faunal communities from highly degraded environmental DNA (Vernot et al. 2021; Zavala et al. 2021). Collectively, these studies established targeted capture as a promising enrichment strategy but no study has evaluated its application as a practical framework for routine, cross-phylum assessment of freshwater ecosystems. Our study extends these efforts by demonstrating that targeted capture can support operational watershed-scale biomonitoring suitable for generation of bioassessment indices.

Compared with unenriched shotgun metagenomics, the principal advantage of targeted capture is sequencing efficiency. In paired libraries generated from the same environmental DNA extracts, targeted capture increased recovery of COI sequences by a median of approximately 1,760-fold. Consequently, obtaining an equivalent number of informative COI reads through shotgun sequencing alone would require on the order of 176 million sequencing reads for a sample that required only 100,000 reads following enrichment. From a cost perspective, for every dollar of sequencing spent on the products of targeted capture, $1,760 in sequencing would be required on the unenriched shotgun libraries. Given the low abundance of target COI in the unenriched libraries, this quickly makes sequencing unenriched libraries uneconomical for the same degree of detection of target organisms.

One of the greatest long-term advantages of targeted capture is its scalability. Unlike multiplexing metabarcoding PCR reactions, where increasing taxonomic breadth requires additional PCR assays, hybridization panels can be expanded simply by incorporating additional baits into the capture pool. This flexibility extends across both taxa and loci. Future panels could simultaneously target multiple mitochondrial and nuclear markers, providing independent confirmation of taxonomic assignments, improved discrimination among closely related species, and greater robustness where any single locus provides limited resolution. Because expanding a capture panel does not alter the underlying laboratory workflow, targeted capture provides a flexible molecular framework that can evolve with reference databases, new genetic markers, and emerging monitoring priorities without requiring wholesale changes to laboratory methods.

### Comparison with conventional monitoring

Comparison with Colorado’s long-term macroinvertebrate monitoring program provides strong support for the targeted capture approach. Mean arthropod richness recovered from a single eDNA sample was remarkably similar to richness reported from the Colorado Multimetric Index (MMI) program within comparable reaches of Boulder Creek, despite fundamentally different sampling methodologies. Combining data from three independent eDNA samples substantially increased taxonomic recovery, yielding considerably higher cumulative richness than conventional morphological surveys. Rather than indicating inflated richness estimates, this pattern likely reflects more complete characterization of local communities through biological replication (Schulte in press; Gold et al. 2023). Thus, a single eDNA sample produced richness comparable to conventional morphology, while replicate sampling progressively expanded the observed community.

The magnitude of invertebrate richness recovered by targeted capture also compares favorably with the largest conventional datasets. Hughes et al. (2023) analyzed 3,358 benthic macroinvertebrate samples from the U.S. EPA National Rivers and Streams Assessment (NRSA), applying genus-level identification to 300 individuals per kick-net sample. Across the conterminous United States, richness averaged 27.4 genera per site, with regional totals of 357– 484 genera across ecoregions. Targeted capture recovered approximately 56 genera per site in Boulder Creek—roughly twice the NRSA average. By the richness–sampling relationship of Hughes et al., the ∼460 genera detected across the watershed would otherwise require on the order of 400 conventional sites. Because our totals include terrestrial and riparian taxa that kick-net sampling would not recover, these comparisons are approximate. Nonetheless, they indicate that targeted capture recovers local assemblages comparable to conventional biomonitoring while achieving cumulative richness that would likely otherwise demand far greater sampling effort.

Although developing or evaluating bioassessment metrics was not our main objective, capture data readily supported two established macroinvertebrate condition indices, both of which indicated declining water quality from headwater to downstream sites. The first, the EPT/(EPT+Chironomidae) richness proportion, is a recognized indicator of stream condition, including fine-sediment impacts (Zweig and Rabeni 2001). Its downstream decline in Boulder Creek matches large-scale conventional biomonitoring, where modified streams lose disturbance-sensitive Ephemeroptera, Plecoptera, Trichoptera, and Megaloptera and become dominated by tolerant chironomids (Rumschlag et al. 2023; Hughes et al. 2023). The second drew on the increaser and decreaser taxa Colorado uses to distinguish pollution-tolerant from pollution-intolerant assemblages (Colorado Department of Public Health and Environment 2017). Pooling designations across the state’s three biotype regions, relative read abundances of decreaser taxa fell downstream while those of increaser taxa rose, with the decreaser proportion declining from near 1.0 in the forested headwaters to near 0 in the agricultural plains. That a narrow, richness-based index and a broader, read-abundance-based index independently recovered the same downstream loss of sensitive taxa reinforces the ecological signal and shows that capture preserves the compositional information these indices depend on. More broadly, deriving established metrics directly from capture data suggests that a range of existing indices could be computed from a single eDNA dataset without additional laboratory work.

### Future directions for targeted capture bioassessment

Several opportunities remain to improve the performance of targeted capture. Increasing the quantity of target DNA entering and retained through hybridization, together with continued optimization of bait design, hybridization chemistry, and sequencing depth, should improve contig assembly and detection sensitivity. Although taxonomy-free assessment is one approach to bioassessment (Cordier et al. 2018), expanding regional reference databases is equally important. Because assignment depends on reference completeness, taxa lacking close references can be confidently misassigned to related but biogeographically implausible genera—as we observed for several mollusc and annelid detections—so broader reference libraries will improve accuracy and reduce reliance on conservative filtering. Targeted capture can also produce longer sequences (*e.g*. full length COI barcodes), which can also be potentially useful in improving reference databases. Coupled with standardized bioinformatic methods for assembling and reporting variable-length contigs, these advances provide a pathway toward increasingly comprehensive and reproducible molecular assessment of aquatic biodiversity.

Realizing this potential will require methodological development on several fronts. Existing bioassessment indices were calibrated to conventional sampling and taxonomic resolution and may need recalibration for molecular datasets that routinely recover greater richness and broader taxonomic coverage (Hughes et al. 2023), especially considering that we ignored richness below the level of genus for this study. The continued accumulation of taxa with replicate sampling further argues for effort-response curves that estimate sampling completeness and asymptotic richness for major taxonomic groups, providing a standardized basis for comparison among sites and sampling efforts (Schulte in press). Quantitative molecular metrics are an equally important opportunity. Although target DNA quantity need not track organism abundance or biomass, owing to species-specific differences in DNA production, transport, and persistence (Yates, Fraser, and Derry 2019; Thalinger et al. 2021), it nonetheless carries ecologically meaningful information about the relative distribution of taxa in space and time. Calibrating DNA quantity against abundance or biomass will require work across taxa and conditions, as will the use of spike-ins to better standardize quantification (Stoeckle, Ausubel, and Coogan 2024), but the variation in recovered read abundances across Boulder Creek and among species suggests that capture preserves ecologically meaningful quantitative differences.

Although richness will remain central to freshwater bioassessment, the broader taxonomic coverage, finer species-level identification, and quantitative signal afforded by capture create opportunities to develop new indices with greater sensitivity and diagnostic power than conventional approaches allow. Beyond improving detection of traditional indicator groups, cross-phylum assessment may better discriminate among stressors rather than reporting only overall condition. Capture will not replace all molecular approaches: for abundant, readily amplified groups such as diatoms, metabarcoding will likely remain simpler and less expensive. Its greatest value lies with diverse assemblages in which target DNA is rare, primer design is challenging, or many taxa must be assessed at once. As bait panels expand across taxa, loci, and regions, targeted capture offers a practical framework for standardized, cross-phylum molecular bioassessment that can grow with reference databases and evolving monitoring needs.

## Acknowledgements

No outside funding was used for this research. Jonah Ventures is a for-profit entity.

## Supplementary Tables

**Table S1.** Site Information. Sampling site metadata: reach category, coordinates, and elevation.

| Category | Site | Latitude | Longitude | Elevation (m) |
| --- | --- | --- | --- | --- |
| Upper Canyon | Lost Lake | 39.9539 | -105.6198 | 2943.4 |
| Upper Canyon | Hessie | 39.9545 | -105.6030 | 2757.1 |
| Upper Canyon | Eldora | 39.9491 | -105.5702 | 2645.2 |
| Upper Canyon | Nederland | 39.9614 | -105.5160 | 2539.9 |
| Lower Canyon | Below Barker | 39.9713 | -105.4733 | 2428.5 |
| Lower Canyon | Castle Rock | 39.9804 | -105.4547 | 2365.5 |
| Lower Canyon | Pulloff | 39.9894 | -105.4276 | 2304.0 |
| Lower Canyon | Eagle Rock | 40.0011 | -105.3894 | 2018.8 |
| Lower Canyon | Fourmile Canyon | 40.0149 | -105.3252 | 1752.8 |
| Boulder | Eben G. Fine | 40.0135 | -105.2947 | 1648.4 |
| Boulder | Boulder High School | 40.0117 | -105.2720 | 1624.6 |
| Boulder | Scott Carpenter | 40.0109 | -105.2548 | 1610.2 |
| Boulder | 55th | 40.0258 | -105.2271 | 1585.6 |
| Boulder | N 75th | 40.0514 | -105.1788 | 1560.1 |
| Plains Reach | N 95th | 40.0504 | -105.1443 | 1544.4 |
| Plains Reach | N155th | 40.0698 | -105.0831 | 1520.2 |
| Plains Reach | CR 16 1/2 | 40.1093 | -105.0369 | 1498.6 |
| Plains Reach | St. Vrain | 40.1704 | -104.9916 | 1474.8 |

**Table S2.**
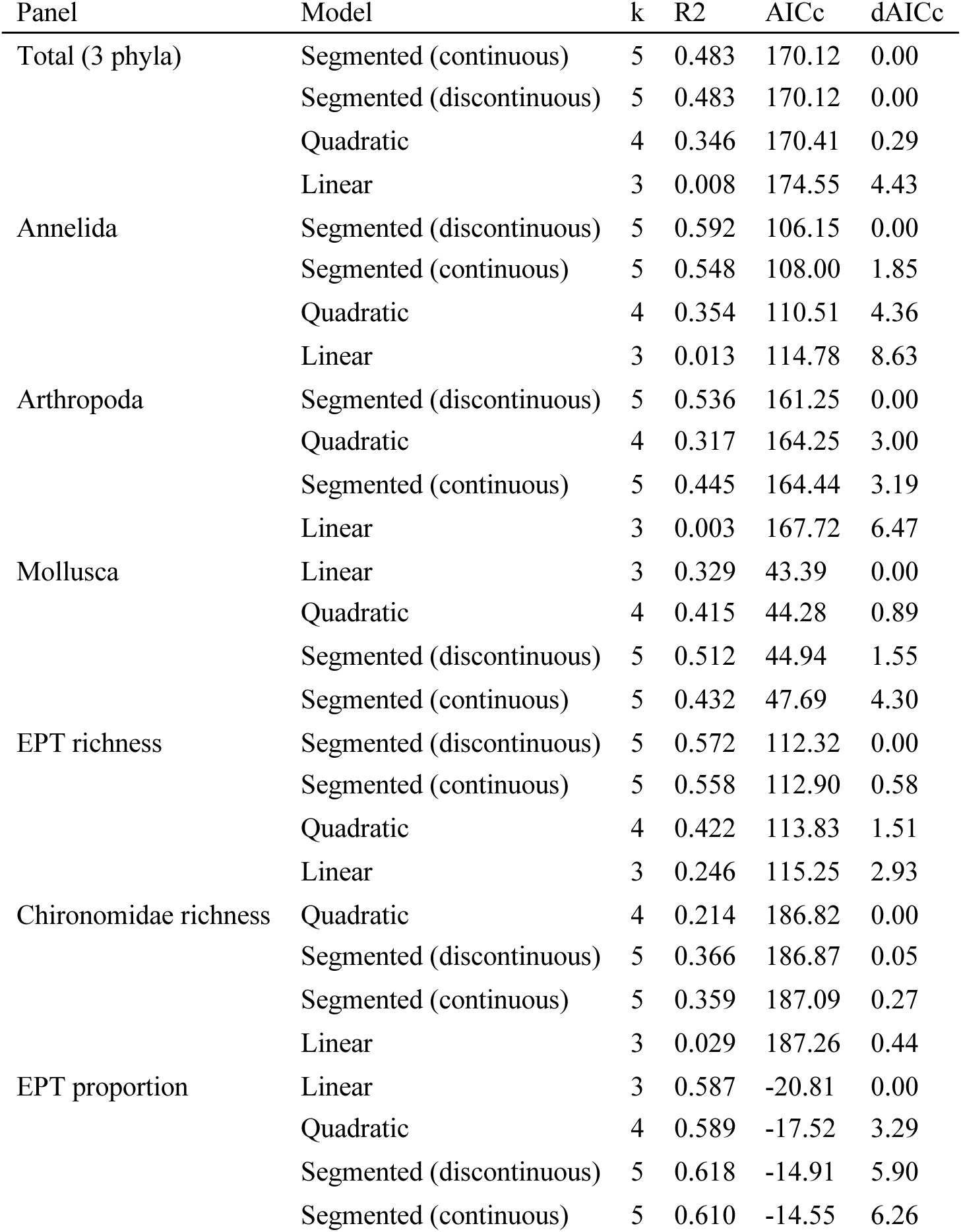
Curve-fit Model Comparison for Richness Metrics vs Distance. Comparison of four candidate models relating mean invertebrate contig richness to river distance for each Figure 5 panel: linear, quadratic, continuous segmented (breakpoint with connected segments), and discontinuous segmented. Models are scored by coefficient of determination (R^2^) and small-sample-corrected Akaike Information Criterion (AICc); *k* is the number of parameters (including residual variance) and ΔAICc is the difference from the best (lowest-AICc) model within each panel.

| Panel | Model | k | R2 | AICc | dAICc |
| --- | --- | --- | --- | --- | --- |
| Total (3 phyla) | Segmented (continuous) | 5 | 0.483 | 170.12 | 0.00 |
|  | Segmented (discontinuous) | 5 | 0.483 | 170.12 | 0.00 |
|  | Quadratic | 4 | 0.346 | 170.41 | 0.29 |
|  | Linear | 3 | 0.008 | 174.55 | 4.43 |
| Annelida | Segmented (discontinuous) | 5 | 0.592 | 106.15 | 0.00 |
|  | Segmented (continuous) | 5 | 0.548 | 108.00 | 1.85 |
|  | Quadratic | 4 | 0.354 | 110.51 | 4.36 |
|  | Linear | 3 | 0.013 | 114.78 | 8.63 |
| Arthropoda | Segmented (discontinuous) | 5 | 0.536 | 161.25 | 0.00 |
|  | Quadratic | 4 | 0.317 | 164.25 | 3.00 |
|  | Segmented (continuous) | 5 | 0.445 | 164.44 | 3.19 |
|  | Linear | 3 | 0.003 | 167.72 | 6.47 |
| Mollusca | Linear | 3 | 0.329 | 43.39 | 0.00 |
|  | Quadratic | 4 | 0.415 | 44.28 | 0.89 |
|  | Segmented (discontinuous) | 5 | 0.512 | 44.94 | 1.55 |
|  | Segmented (continuous) | 5 | 0.432 | 47.69 | 4.30 |
| EPT richness | Segmented (discontinuous) | 5 | 0.572 | 112.32 | 0.00 |
|  | Segmented (continuous) | 5 | 0.558 | 112.90 | 0.58 |
|  | Quadratic | 4 | 0.422 | 113.83 | 1.51 |
|  | Linear | 3 | 0.246 | 115.25 | 2.93 |
| Chironomidae richness | Quadratic | 4 | 0.214 | 186.82 | 0.00 |
|  | Segmented (discontinuous) | 5 | 0.366 | 186.87 | 0.05 |
|  | Segmented (continuous) | 5 | 0.359 | 187.09 | 0.27 |
|  | Linear | 3 | 0.029 | 187.26 | 0.44 |
| EPT proportion | Linear | 3 | 0.587 | -20.81 | 0.00 |
|  | Quadratic | 4 | 0.589 | -17.52 | 3.29 |
|  | Segmented (discontinuous) | 5 | 0.618 | -14.91 | 5.90 |
|  | Segmented (continuous) | 5 | 0.610 | -14.55 | 6.26 |

**Table S3.** Sequence Length Distribution (Folmer-trimmed & Untrimmed). Distribution of sequence lengths binned across the three main invertebrate phyla (Arthropoda, Annelida, Mollusca), shown for both Folmer-trimmed barcodes and untrimmed assembled contigs. Counts and within-group percentages are given for each length bin.

| Length<br>bin (bp) | Invert<br>Folmer<br>(n) | Invert<br>Folmer<br>(%) | Invert<br>untrimmed<br>(n) | Invert<br>untrimmed<br>(%) |
| --- | --- | --- | --- | --- |
| <200 | 283 | 11.0 | 0 | 0.0 |
| 200-249 | 297 | 11.5 | 69 | 2.7 |
| 250-299 | 297 | 11.5 | 69 | 2.7 |
| 300-349 | 421 | 16.3 | 436 | 16.9 |
| 350-399 | 295 | 11.4 | 385 | 14.9 |
| 400-449 | 220 | 8.5 | 311 | 12.0 |
| 450-499 | 140 | 5.4 | 233 | 9.0 |
| 500-549 | 140 | 5.4 | 176 | 6.8 |
| 550-599 | 178 | 6.9 | 149 | 5.8 |
| 600-649 | 146 | 5.7 | 120 | 4.6 |
| ≥650 | 166 | 6.4 | 635 | 24.6 |

**Table S4.** Genus Accumulation Across Samples. Mean number of invertebrate genera (Annelida, Arthropoda, Mollusca) accumulated as the three within-site samples are pooled, averaged across the 18 sites. ‘New genera’ is the mean number of additional genera contributed by each successive sample; ‘% of total’ expresses the running mean as a percentage of the three-sample total.

| Samples pooled | Mean genera | SD | SE | New genera | % of total |
| --- | --- | --- | --- | --- | --- |
| 1 | 27.5 | 10.8 | 2.5 | 27.5 | 48.8 |
| 2 | 43.6 | 16.4 | 3.9 | 16.1 | 77.4 |
| 3 | 56.4 | 20.6 | 4.9 | 12.7 | 100 |

**Table S5.**
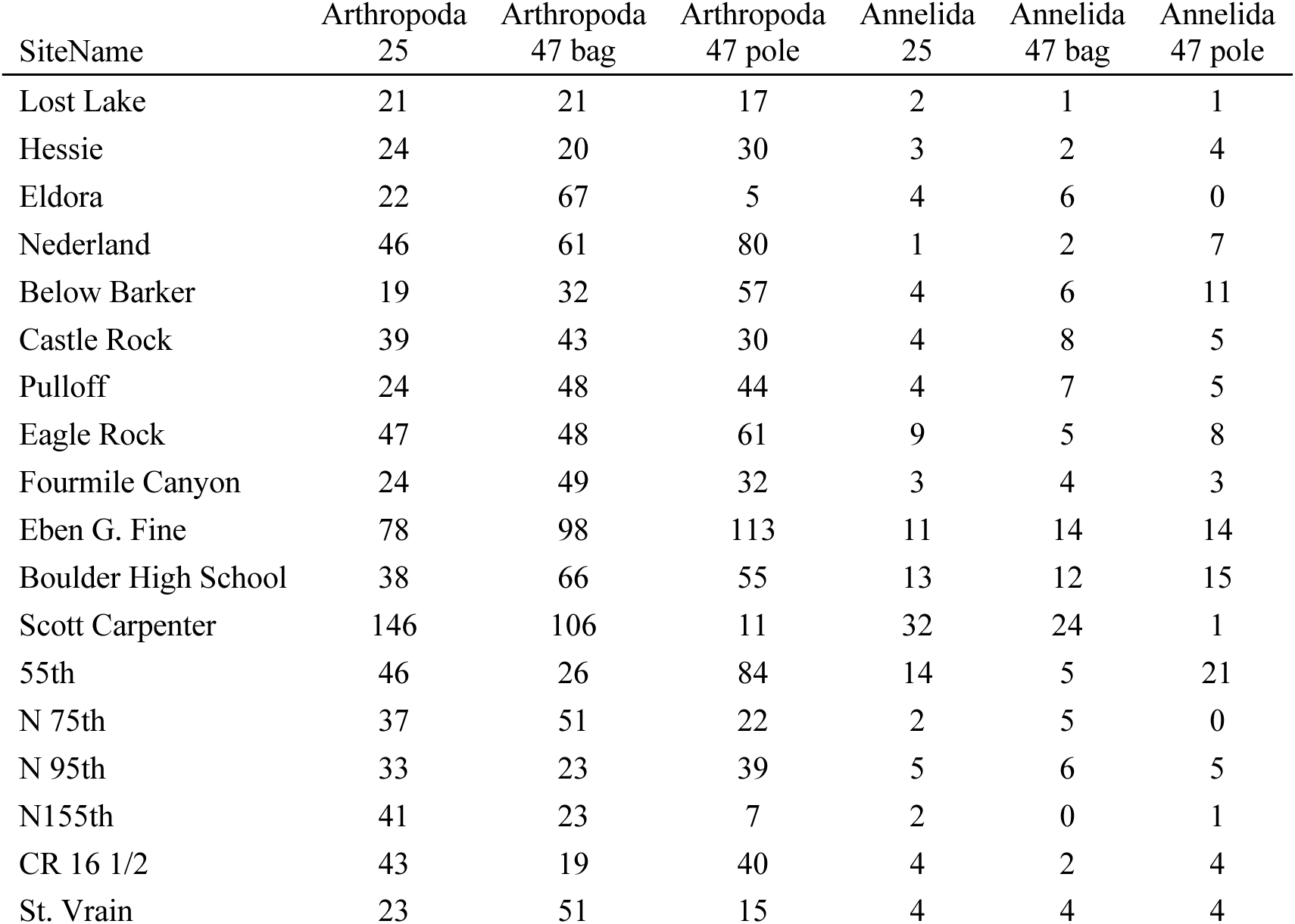
Site-level Arthropod & Annelid Richness by Sample Type. Included are data for each of the three sampling types (25-mm filter, 47-mm filter with bag filtering, and 47-mm filter with direct filtering)

| SiteName | Arthropoda<br>25 | Arthropoda<br>47 bag | Arthropoda<br>47 pole | Annelida<br>25 | Annelida<br>47 bag | Annelida<br>47 pole |
| --- | --- | --- | --- | --- | --- | --- |
| Lost Lake | 21 | 21 | 17 | 2 | 1 | 1 |
| Hessie | 24 | 20 | 30 | 3 | 2 | 4 |
| Eldora | 22 | 67 | 5 | 4 | 6 | 0 |
| Nederland | 46 | 61 | 80 | 1 | 2 | 7 |
| Below Barker | 19 | 32 | 57 | 4 | 6 | 11 |
| Castle Rock | 39 | 43 | 30 | 4 | 8 | 5 |
| Pulloff | 24 | 48 | 44 | 4 | 7 | 5 |
| Eagle Rock | 47 | 48 | 61 | 9 | 5 | 8 |
| Fourmile Canyon | 24 | 49 | 32 | 3 | 4 | 3 |
| Eben G. Fine | 78 | 98 | 113 | 11 | 14 | 14 |
| Boulder High School | 38 | 66 | 55 | 13 | 12 | 15 |
| Scott Carpenter | 146 | 106 | 11 | 32 | 24 | 1 |
| 55th | 46 | 26 | 84 | 14 | 5 | 21 |
| N 75th | 37 | 51 | 22 | 2 | 5 | 0 |
| N 95th | 33 | 23 | 39 | 5 | 6 | 5 |
| N155th | 41 | 23 | 7 | 2 | 0 | 1 |
| CR 16 1/2 | 43 | 19 | 40 | 4 | 2 | 4 |
| St. Vrain | 23 | 51 | 15 | 4 | 4 | 4 |

**Table S6.** Invertebrate Richness Summary by Sample Type.

|  | Annelida |  | Arthropoda |  | Volume (mL) |
| --- | --- | --- | --- | --- | --- |
|  | Richness (mean<br>± SD) | Range | Richness (mean<br>± SD) | Range |  |
| 25mm disc | 6.7 ± 7.4<sup>a</sup> | 1–32 | 41.7 ± 29.7<sup>a</sup> | 19–146 | 297 ± 158 |
| Unio baleen | 6.6 ± 5.6<sup>a</sup> | 1–24 | 47.3 ± 25.6<sup>a</sup> | 19–106 | 752 ± 345 |
| Unio pole | 6.8 ± 5.7<sup>a</sup> | 1–21 | 41.2 ± 29.4<sup>a</sup> | 5–113 | 656 ± 453 |

Superscript letters denote post-hoc groupings (Kruskal-Wallis followed by pairwise Wilcoxon tests, Benjamini-Hochberg adjusted). Sample types sharing a letter do not differ significantly (alpha = 0.05). Omnibus Kruskal-Wallis: Annelida $p = 0.694$ ; Arthropoda $p = 0.485$ .

**Table S7.**
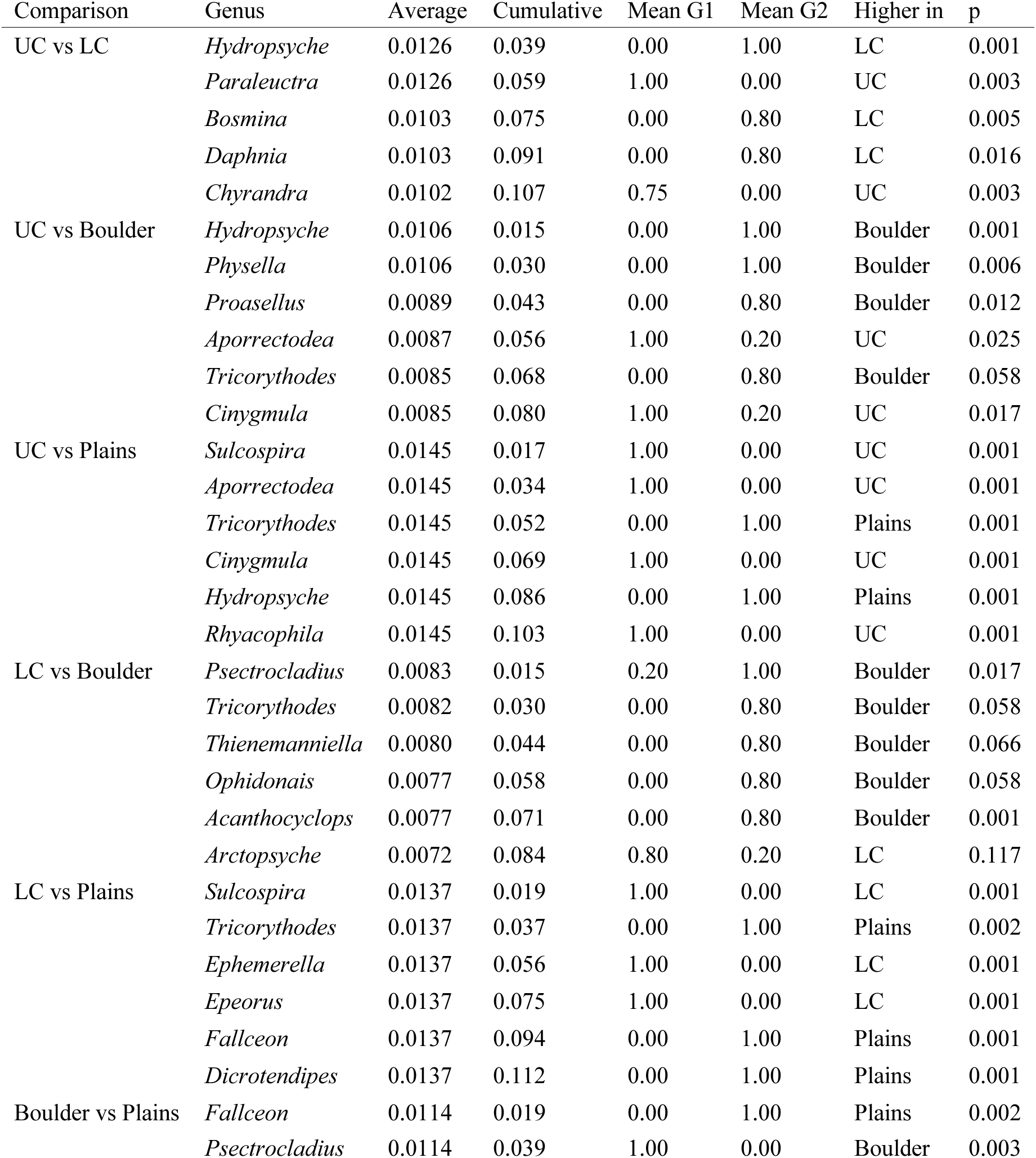
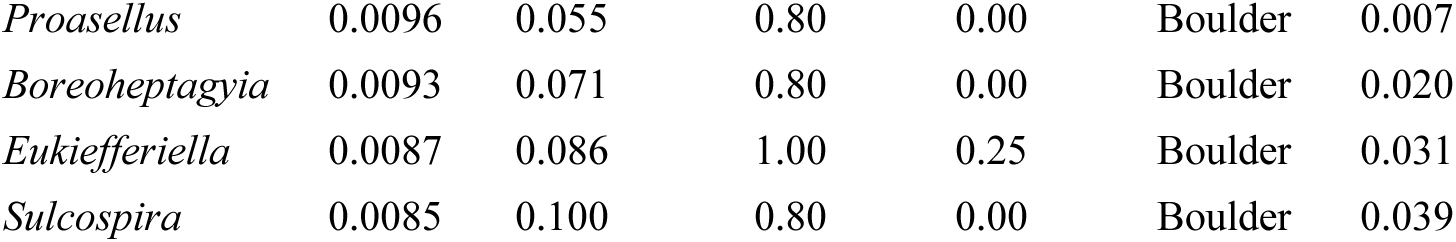
SIMPER: Genera Driving Compositional Differences Among Reaches. Analysis restricted to the three main invertebrate phyla (Annelida, Arthropoda, Mollusca). Top genera (up to six per comparison) contributing to average between-reach compositional dissimilarity (genus-level Jaccard). ‘Average’ is the mean contribution to overall dissimilarity; ‘Cumulative’ is the running proportion of dissimilarity explained. ‘Mean (group 1)’ and ‘Mean (group 2)’ are the genus occurrence frequencies (proportion of sites detected, 0–1) in the first and second reach of the comparison, respectively (the matrix is presence/absence, so these are detection frequencies rather than read abundances); ‘Higher in’ names the reach where the genus is more frequently detected. *p* from 999 permutations. Reaches are Upper Canyon (UC), Lower Canyon (LC), Boulder, and Plains.

| Comparison | Genus | Average | Cumulative | Mean G1 | Mean G2 | Higher in | p |
| --- | --- | --- | --- | --- | --- | --- | --- |
| UC vs LC | *Hydropsyche* | 0.0126 | 0.039 | 0.00 | 1.00 | LC | 0.001 |
|  | *Paraleuctra* | 0.0126 | 0.059 | 1.00 | 0.00 | UC | 0.003 |
|  | *Bosmina* | 0.0103 | 0.075 | 0.00 | 0.80 | LC | 0.005 |
|  | *Daphnia* | 0.0103 | 0.091 | 0.00 | 0.80 | LC | 0.016 |
|  | *Chyrandra* | 0.0102 | 0.107 | 0.75 | 0.00 | UC | 0.003 |
| UC vs Boulder | *Hydropsyche* | 0.0106 | 0.015 | 0.00 | 1.00 | Boulder | 0.001 |
|  | *Physella* | 0.0106 | 0.030 | 0.00 | 1.00 | Boulder | 0.006 |
|  | *Proasellus* | 0.0089 | 0.043 | 0.00 | 0.80 | Boulder | 0.012 |
|  | *Aporrectodea* | 0.0087 | 0.056 | 1.00 | 0.20 | UC | 0.025 |
|  | *Tricorythodes* | 0.0085 | 0.068 | 0.00 | 0.80 | Boulder | 0.058 |
|  | *Cinygmula* | 0.0085 | 0.080 | 1.00 | 0.20 | UC | 0.017 |
| UC vs Plains | *Sulcospira* | 0.0145 | 0.017 | 1.00 | 0.00 | UC | 0.001 |
|  | *Aporrectodea* | 0.0145 | 0.034 | 1.00 | 0.00 | UC | 0.001 |
|  | *Tricorythodes* | 0.0145 | 0.052 | 0.00 | 1.00 | Plains | 0.001 |
|  | *Cinygmula* | 0.0145 | 0.069 | 1.00 | 0.00 | UC | 0.001 |
|  | *Hydropsyche* | 0.0145 | 0.086 | 0.00 | 1.00 | Plains | 0.001 |
|  | *Rhyacophila* | 0.0145 | 0.103 | 1.00 | 0.00 | UC | 0.001 |
| LC vs Boulder | *Psectrocladius* | 0.0083 | 0.015 | 0.20 | 1.00 | Boulder | 0.017 |
|  | *Tricorythodes* | 0.0082 | 0.030 | 0.00 | 0.80 | Boulder | 0.058 |
|  | *Thienemanniella* | 0.0080 | 0.044 | 0.00 | 0.80 | Boulder | 0.066 |
|  | *Ophidonais* | 0.0077 | 0.058 | 0.00 | 0.80 | Boulder | 0.058 |
|  | *Acanthocyclops* | 0.0077 | 0.071 | 0.00 | 0.80 | Boulder | 0.001 |
|  | *Arctopsyche* | 0.0072 | 0.084 | 0.80 | 0.20 | LC | 0.117 |
| LC vs Plains | *Sulcospira* | 0.0137 | 0.019 | 1.00 | 0.00 | LC | 0.001 |
|  | *Tricorythodes* | 0.0137 | 0.037 | 0.00 | 1.00 | Plains | 0.002 |
|  | *Ephemerella* | 0.0137 | 0.056 | 1.00 | 0.00 | LC | 0.001 |
|  | *Epeorus* | 0.0137 | 0.075 | 1.00 | 0.00 | LC | 0.001 |
|  | *Fallceon* | 0.0137 | 0.094 | 0.00 | 1.00 | Plains | 0.001 |
|  | *Dicrotendipes* | 0.0137 | 0.112 | 0.00 | 1.00 | Plains | 0.001 |
| Boulder vs Plains | *Fallceon* | 0.0114 | 0.019 | 0.00 | 1.00 | Plains | 0.002 |
|  | *Psectrocladius* | 0.0114 | 0.039 | 1.00 | 0.00 | Boulder | 0.003 |

|  |  |  |  |  |  |  |
| --- | --- | --- | --- | --- | --- | --- |
| *Proasellus* | 0.0096 | 0.055 | 0.80 | 0.00 | Boulder | 0.007 |
| *Boreoheptagyia* | 0.0093 | 0.071 | 0.80 | 0.00 | Boulder | 0.020 |
| *Eukiefferiella* | 0.0087 | 0.086 | 1.00 | 0.25 | Boulder | 0.031 |
| *Sulcospira* | 0.0085 | 0.100 | 0.80 | 0.00 | Boulder | 0.039 |

**Table S8.**
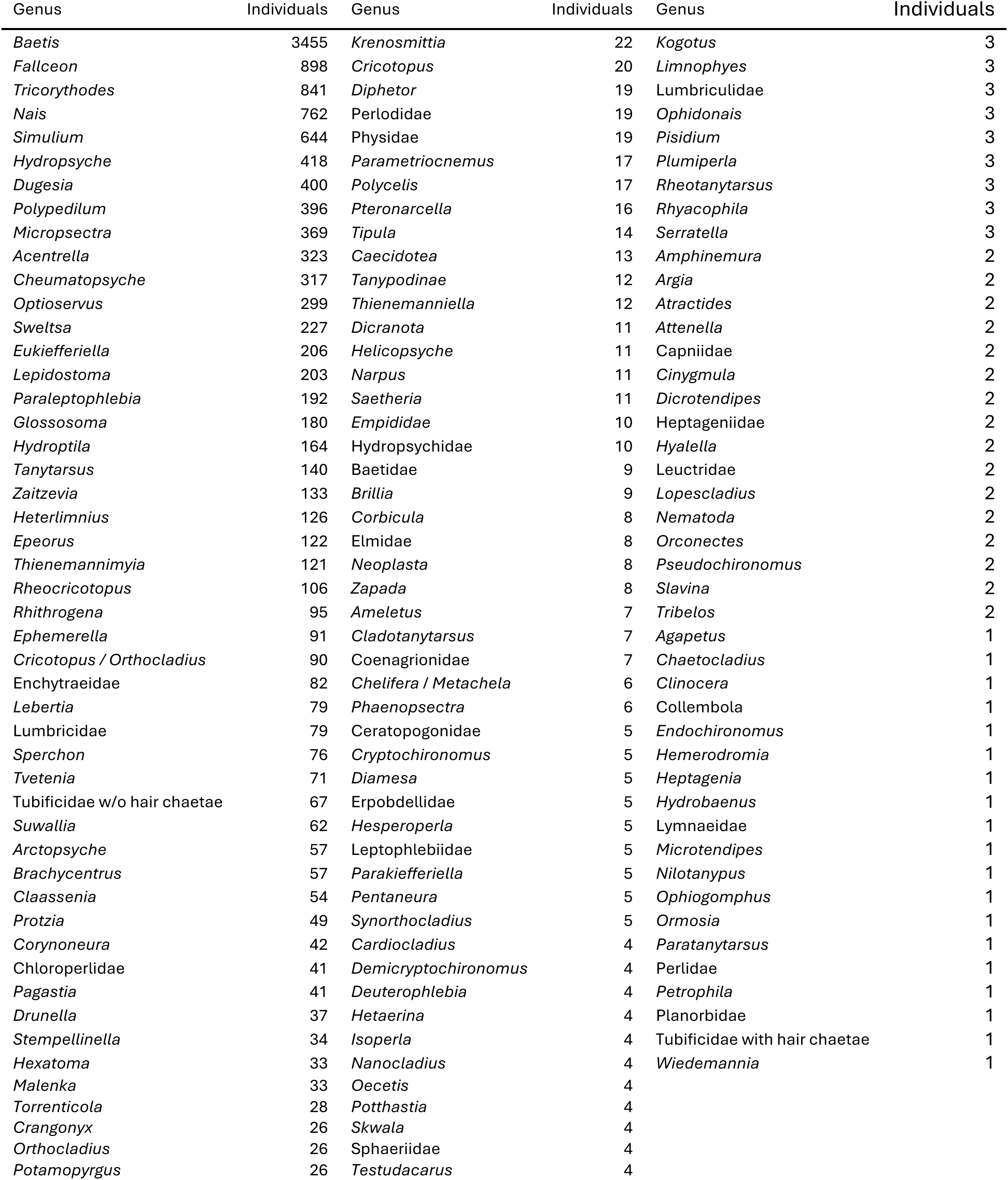
Colorado MMI Total Individuals by Genus. Total number of individuals (Colorado MMI program, CDPHE EDAS) summed by the genus of each FinalID from the ColoradoMMI Data sheet.

**Table S9.** MMI vs eDNA Arthropod Richness by Reach. Comparison of arthropod richness from morphological sampling (Colorado MMI program) and eDNA capture, summarized separately for the Canyon and Boulder-Plains reaches. MMI canyon richness is from the two Middle Boulder Creek stations (Nederland, below Barker; ∼10.0–15.0 km from Lost Lake); the eDNA canyon metric is restricted to the eDNA sites falling between and/or adjacent to those stations (Nederland, Below Barker, Castle Rock, Pulloff). MMI values are counts of morphological taxa; eDNA values are mean contig richness.

| Method | Reach | n | Mean | SD | Median | Min | Max |
| --- | --- | --- | --- | --- | --- | --- | --- |
| MMI arthropod (morphology) | Canyon | 2 | 42.5 | 4.9 | 42.5 | 39.0 | 46.0 |
| eDNA arthropod (capture) | Canyon | 4 | 41.3 | 14.9 | 38.0 | 27.0 | 62.3 |
| MMI arthropod (morphology) | Boulder-Plains | 8 | 31.9 | 10.9 | 30.5 | 18.0 | 51.0 |
| eDNA arthropod (capture) | Boulder-Plains | 9 | 49.4 | 26.1 | 36.7 | 23.7 | 96.3 |

**Table S10.**
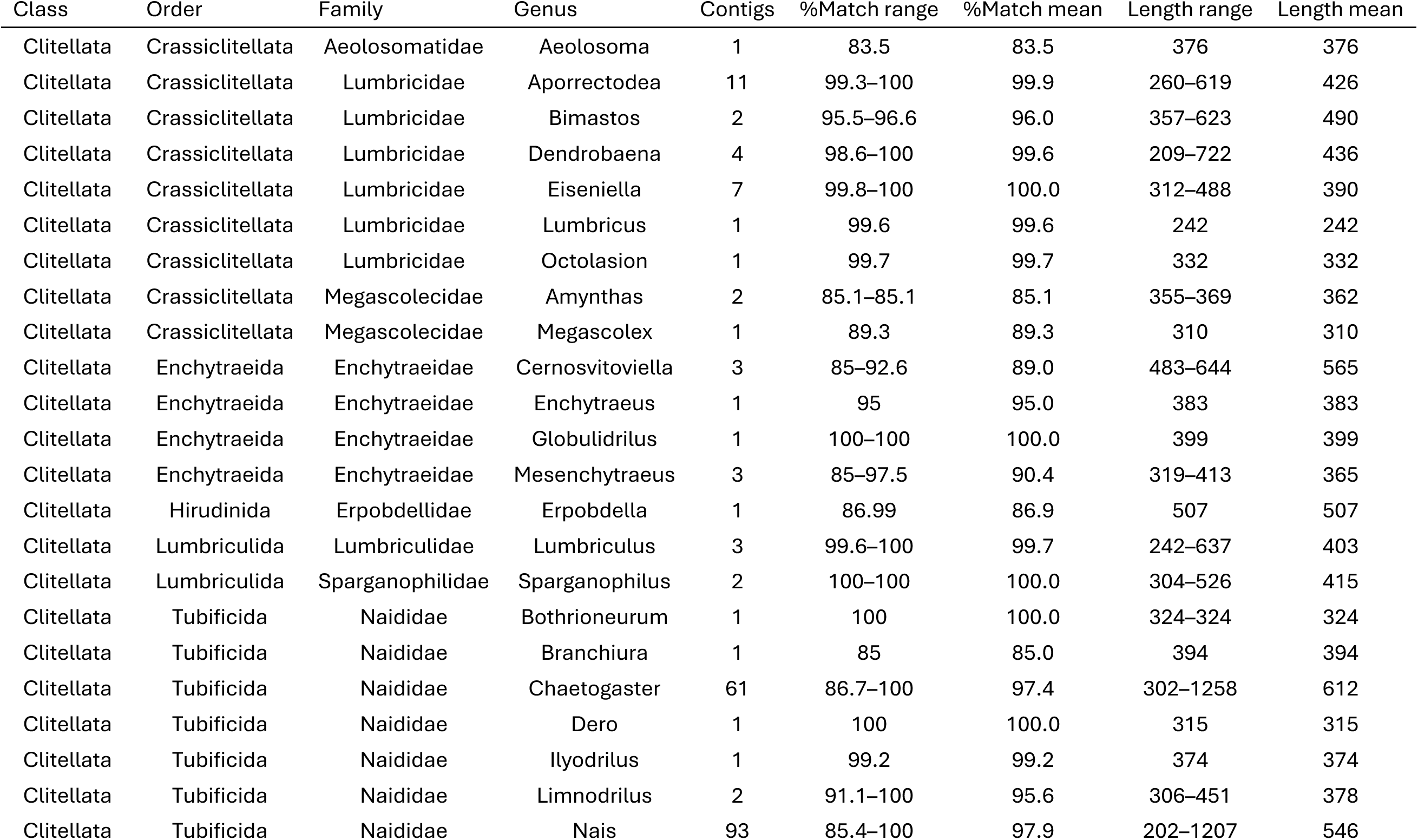
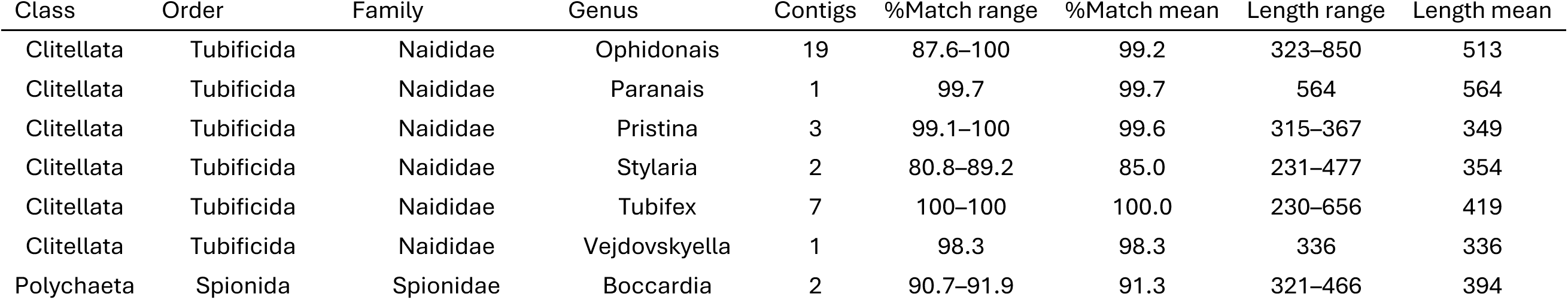
Annelida Taxonomic List. Unique class/order/family/genus list for Annelida Taxonomic List from the cleaned community (trusted genera only). Each genus appears once using its majority-vote class/order/family path; ‘Contigs’ is the number of contigs observed, ‘%Match’ is the genus-supplying percent identity (range and mean), and ‘Length’ is the trimmed contig length (range and mean, bp). Ranks left as NA were never confidently resolved or were dropped as inconsistent with the contig phylum.

**Table S11.**
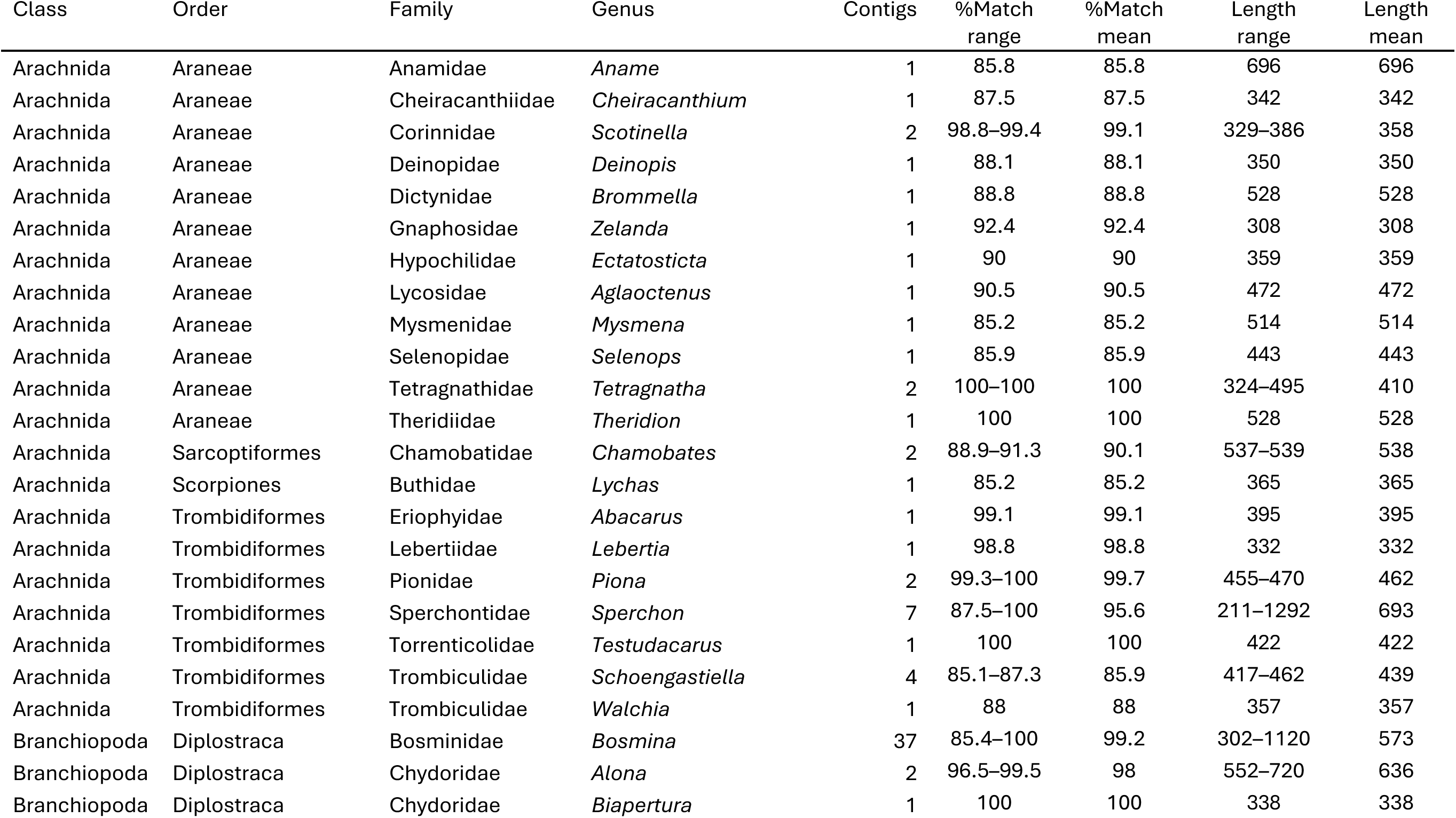
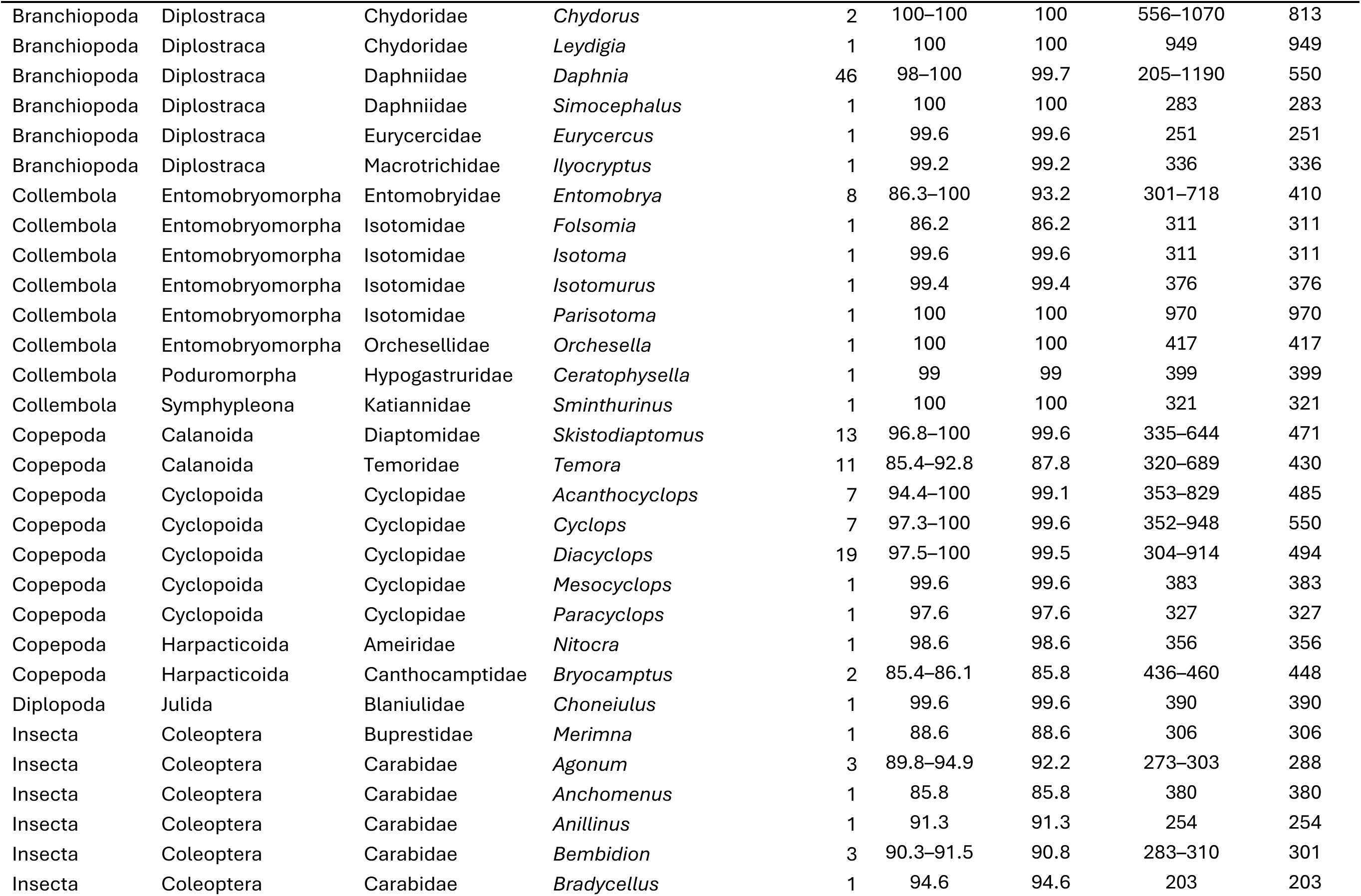
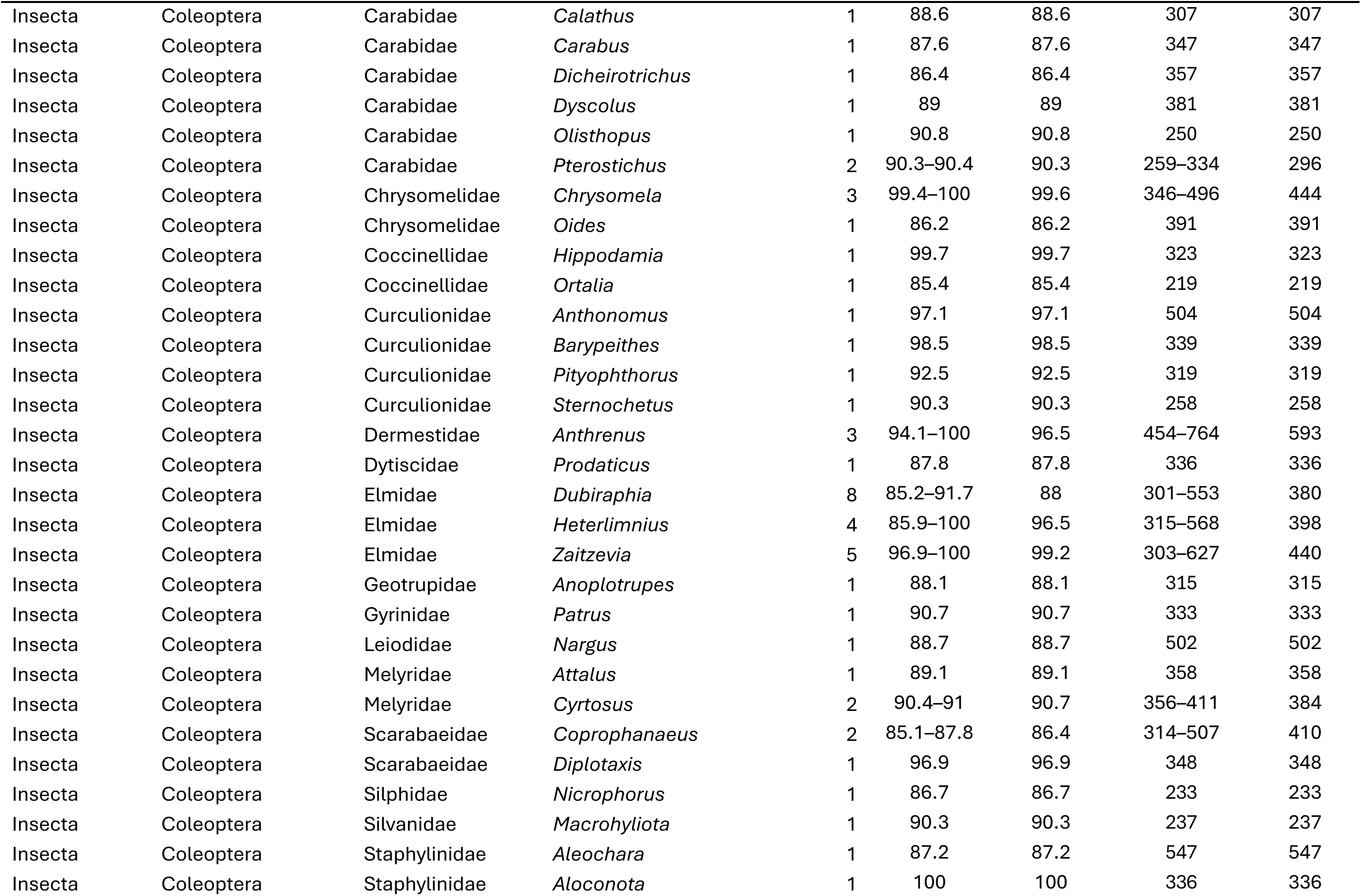
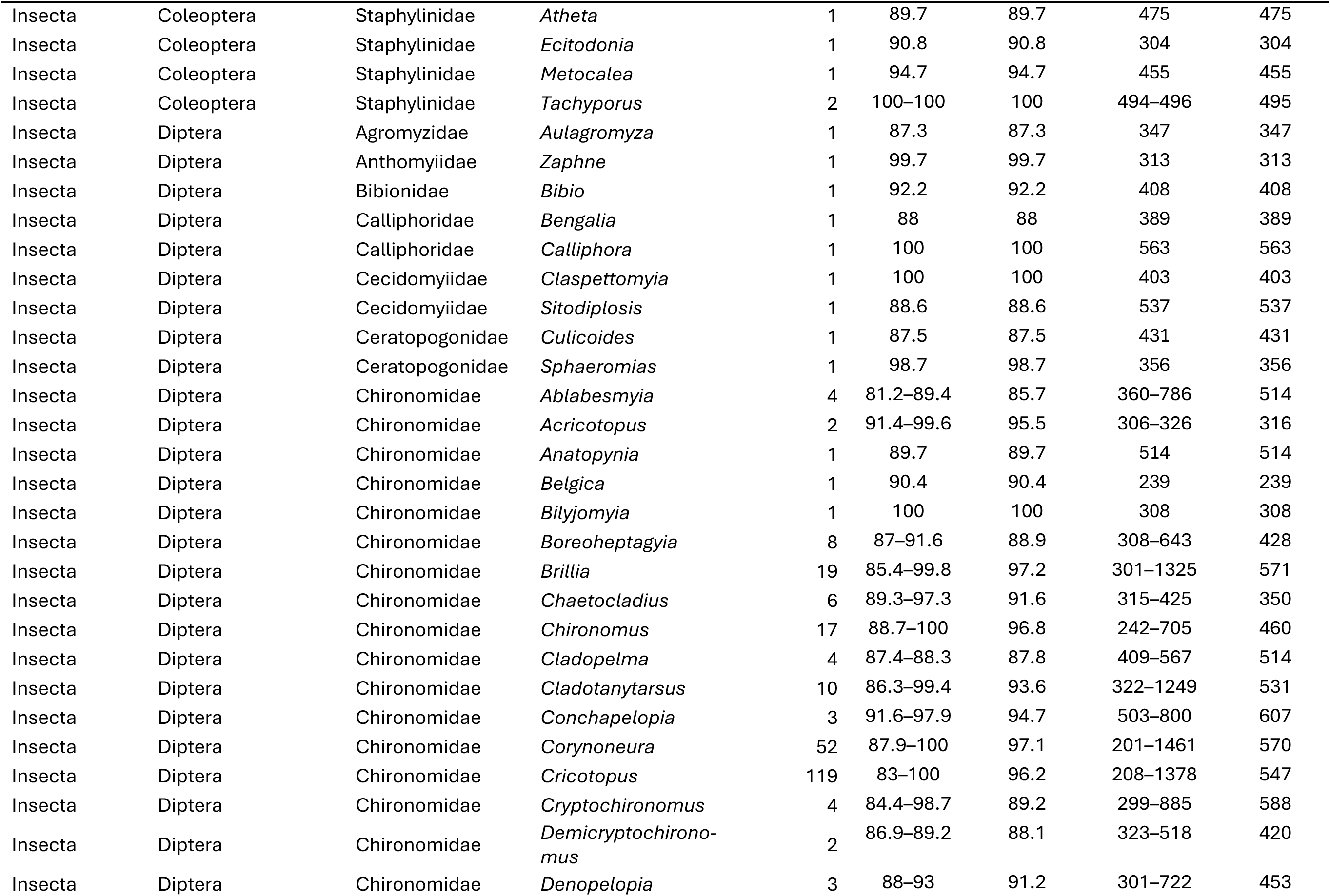
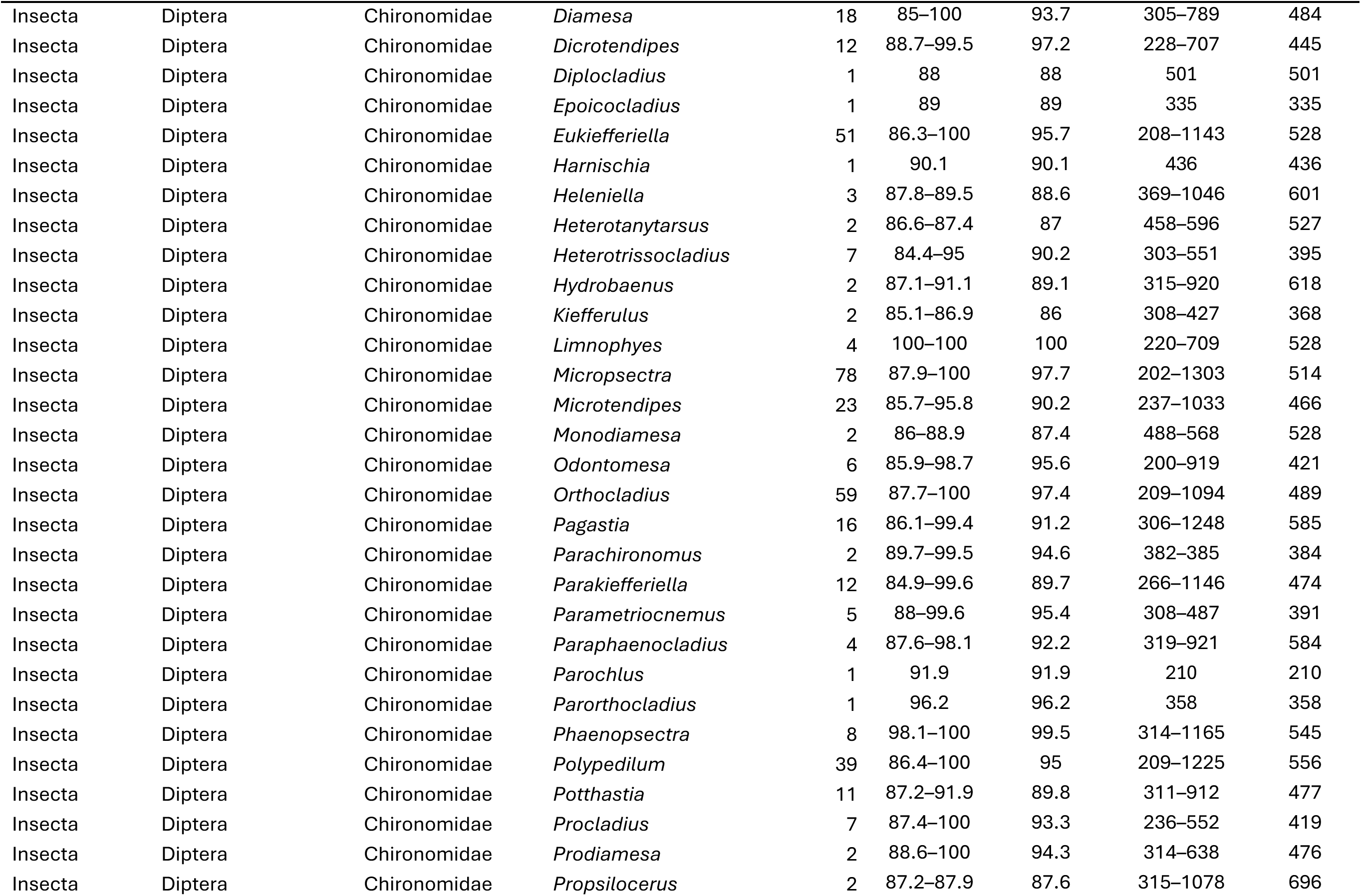
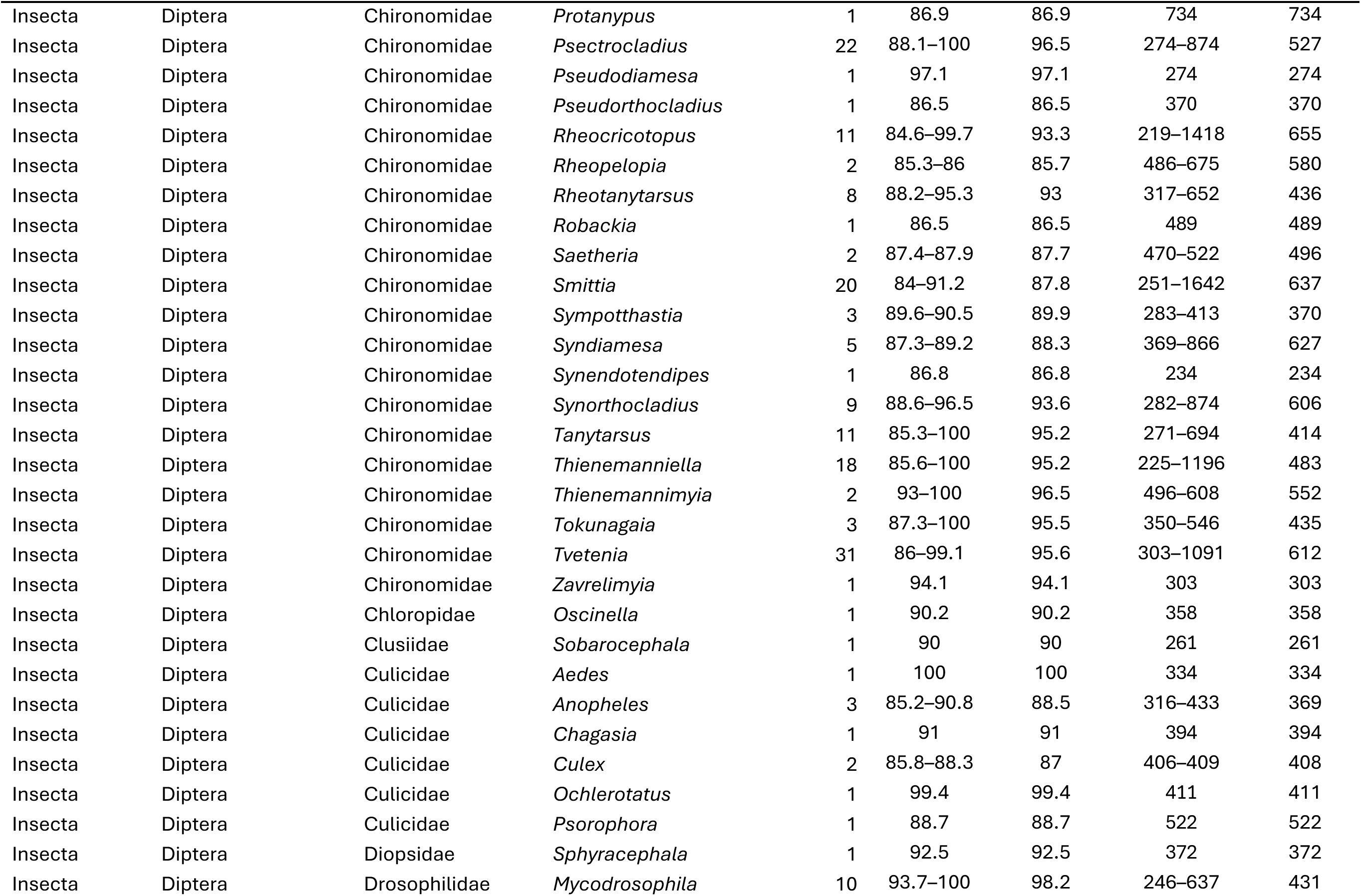
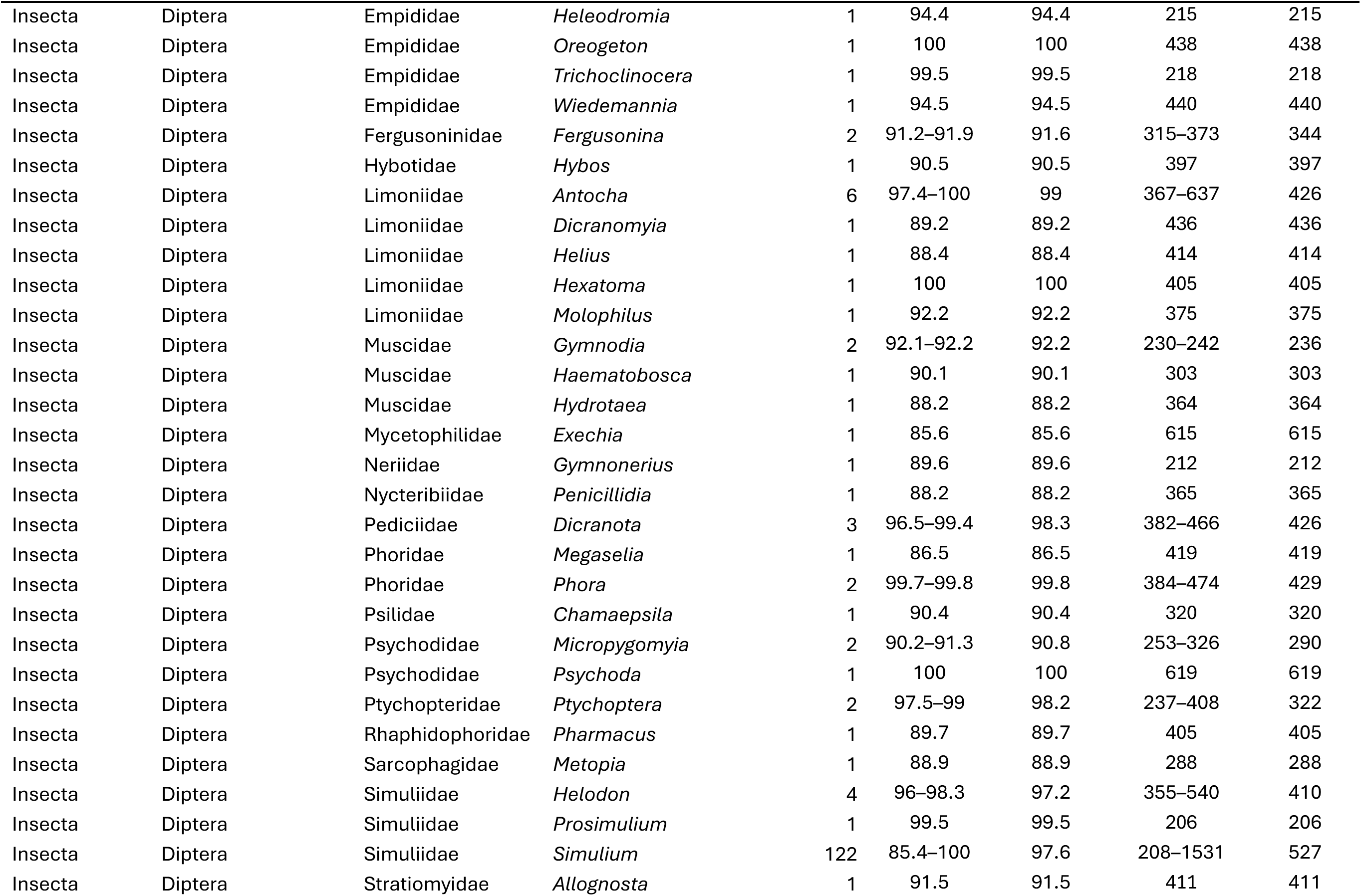
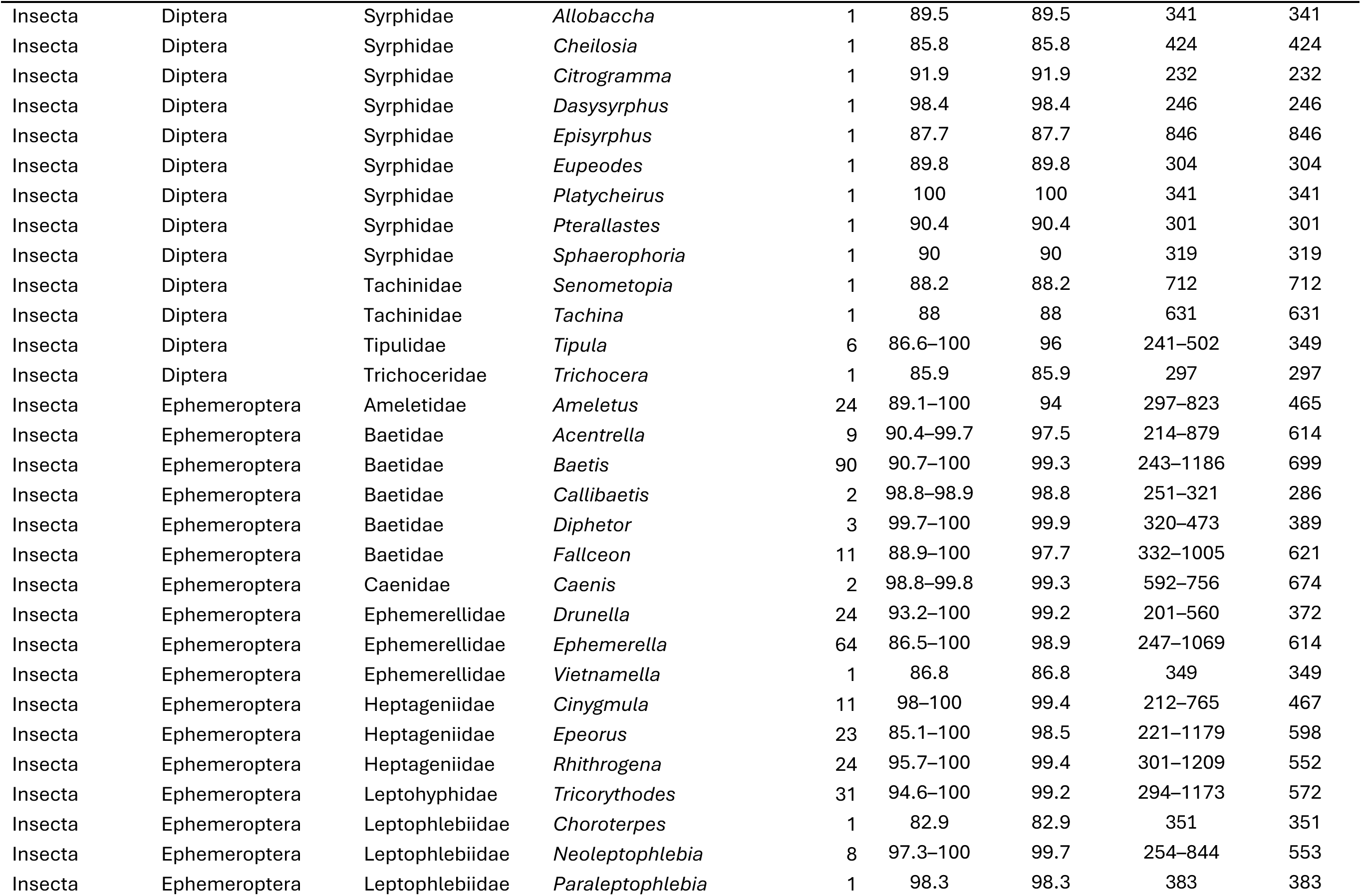
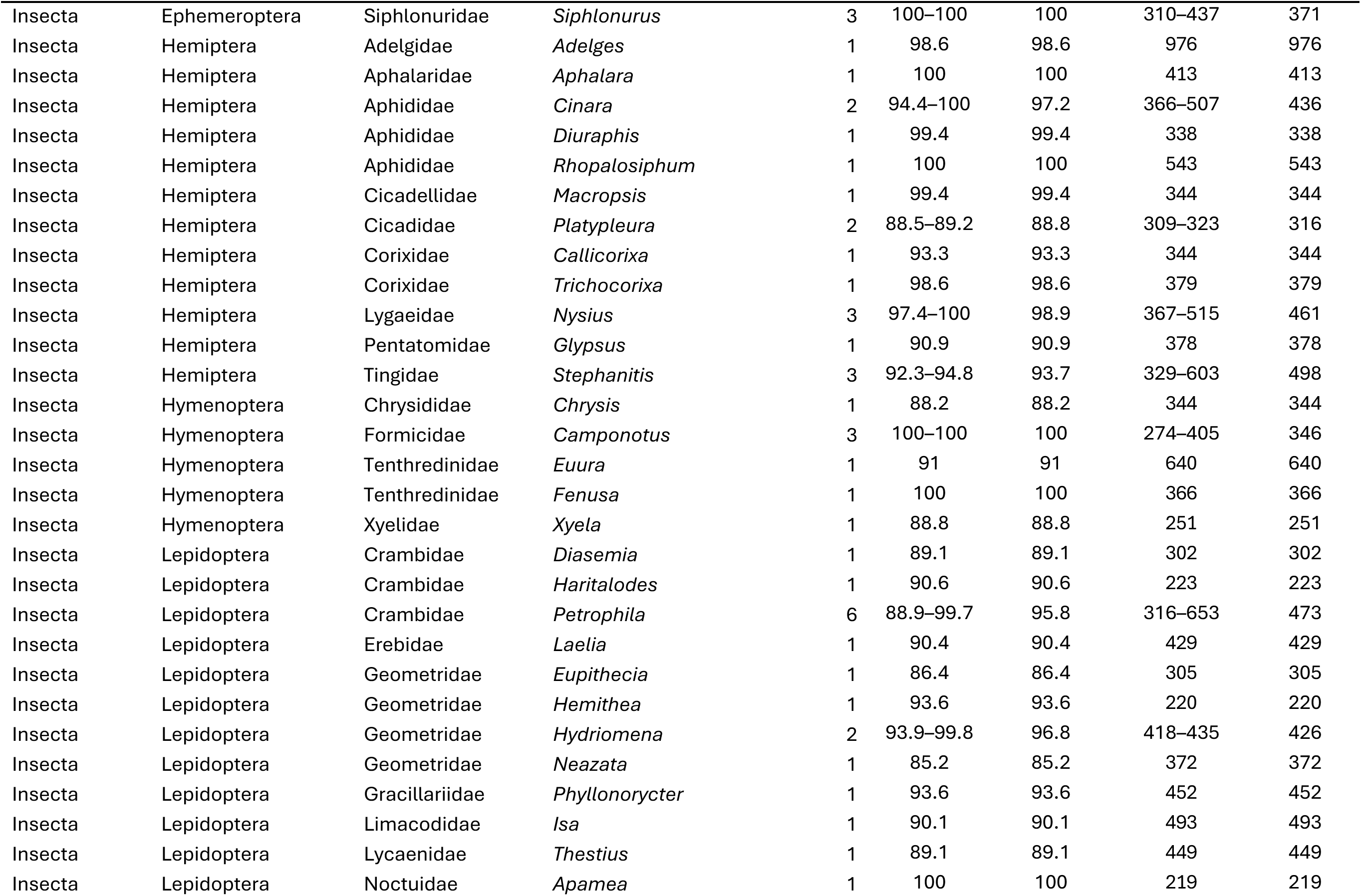
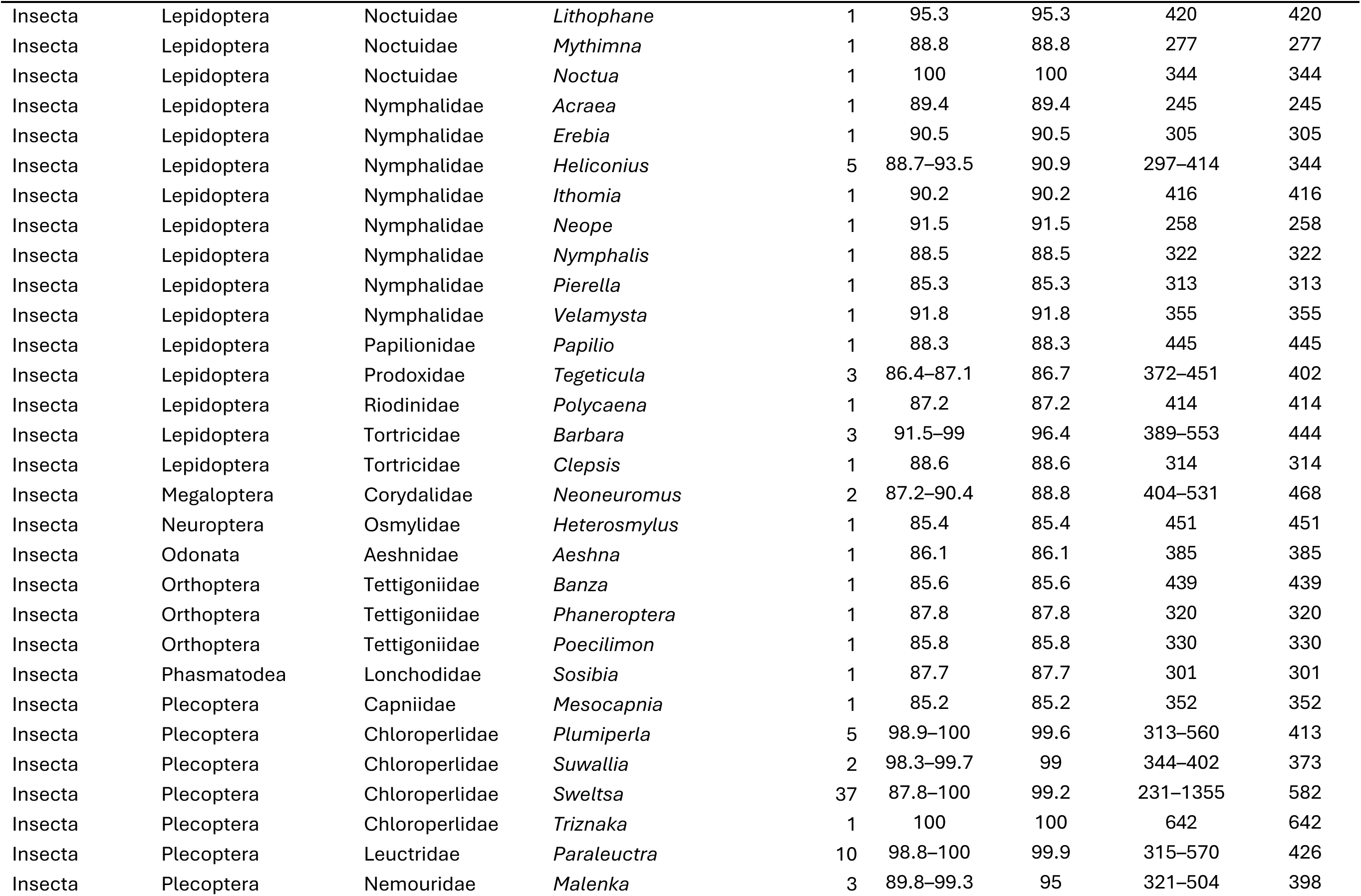
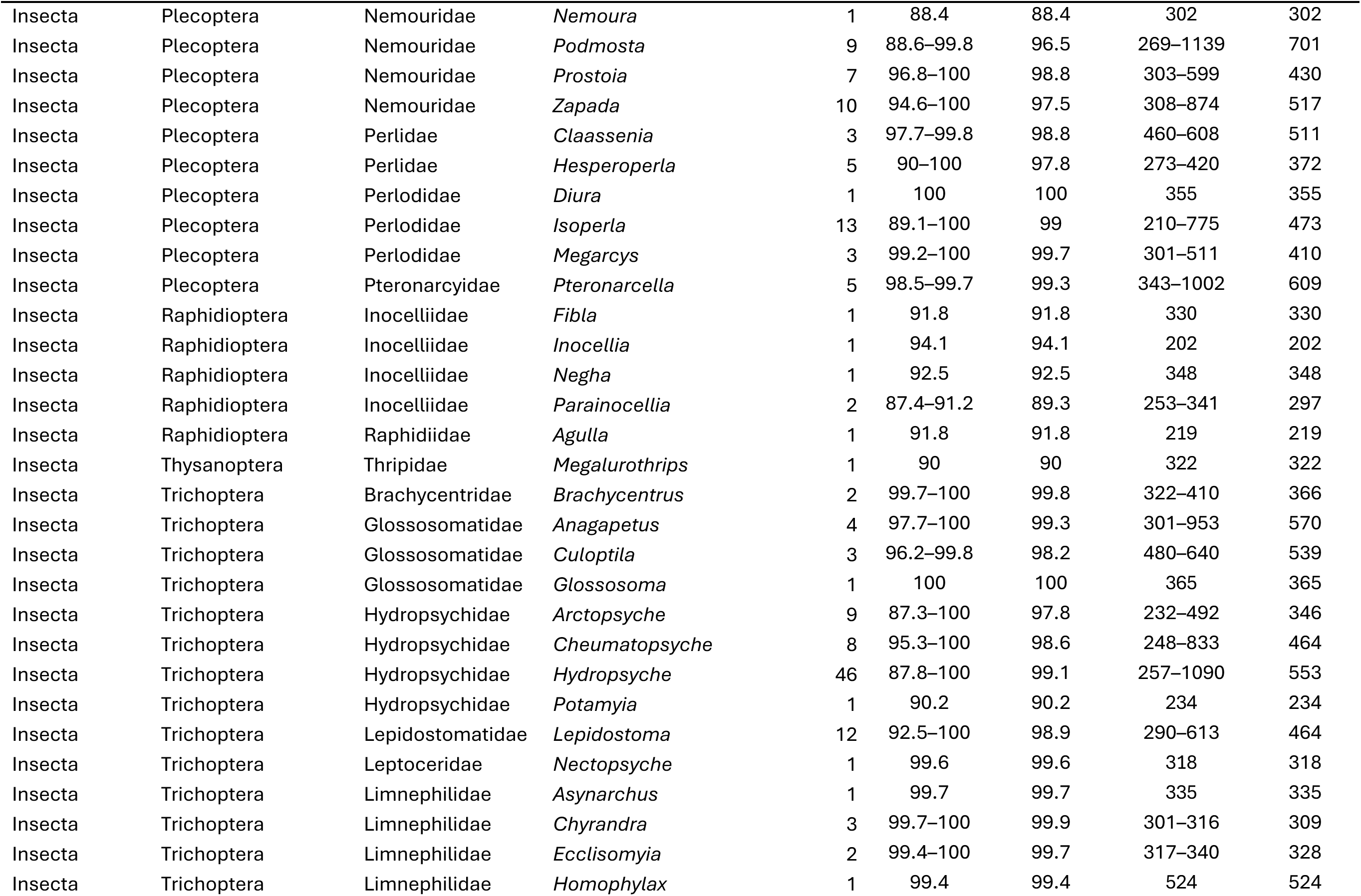
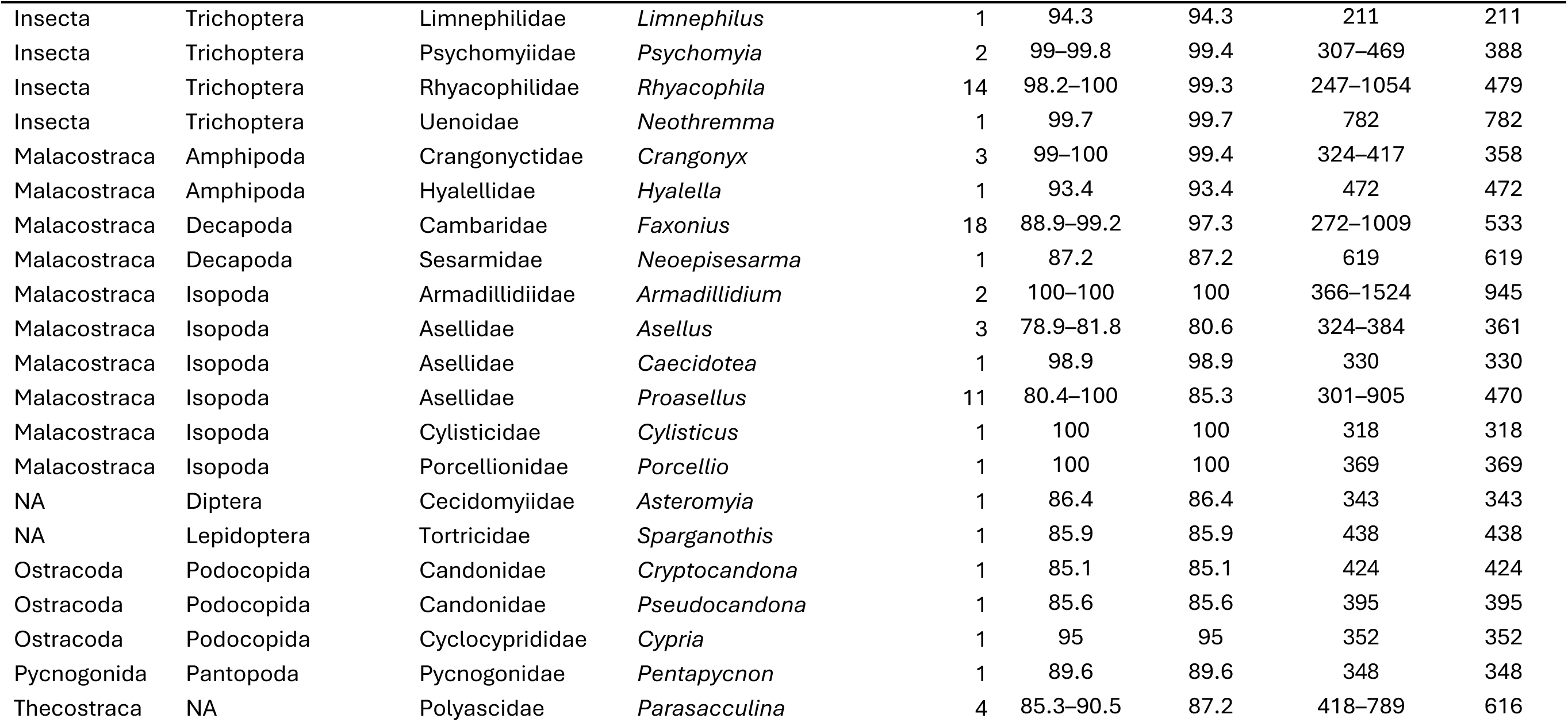
Arthropoda Taxonomic List. Unique class/order/family/genus list for Arthropoda Taxonomic List from the cleaned community (trusted genera only). Each genus appears once using its majority-vote class/order/family path; ‘Contigs’ is the number of contigs observed, ‘%Match’ is the genus-supplying percent identity (range and mean), and ‘Length’ is the trimmed contig length (range and mean, bp). Ranks left as NA were never confidently resolved or were dropped as inconsistent with the contig phylum.

**Table S12.**
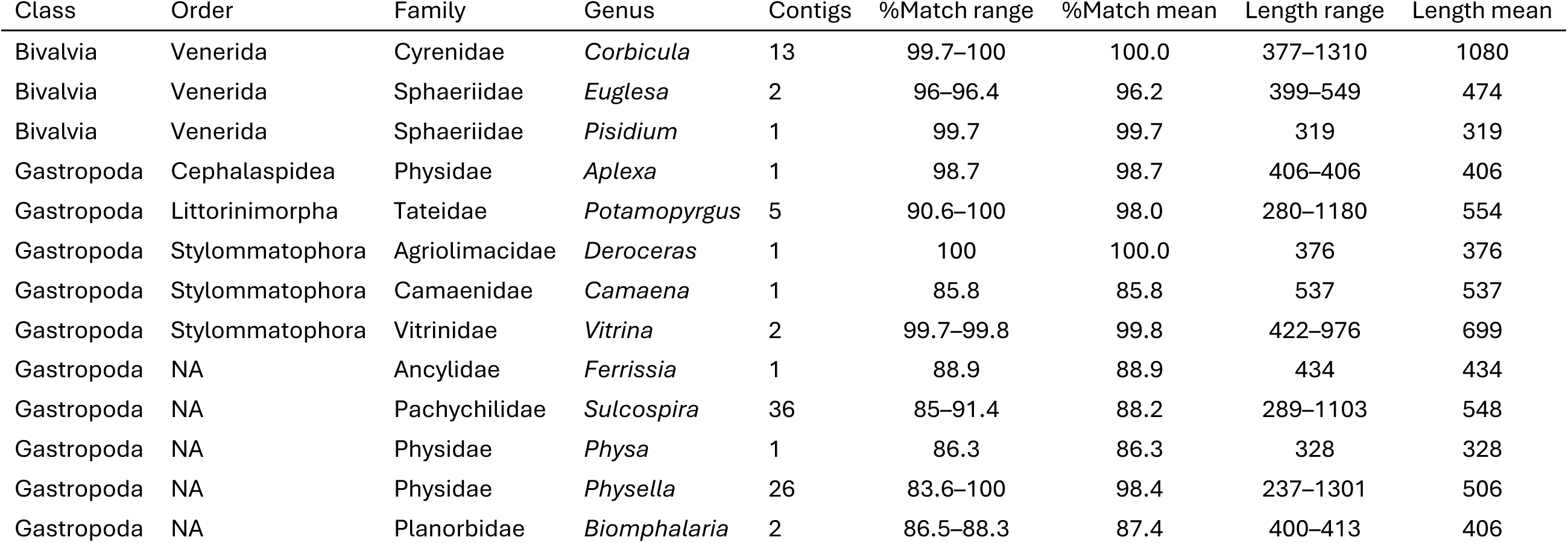
Mollusca Taxonomic List. Unique class/order/family/genus list for Mollusca Taxonomic List from the cleaned community (trusted genera only). Each genus appears once using its majority-vote class/order/family path; ‘Contigs’ is the number of contigs observed, ‘%Match’ is the genus-supplying percent identity (range and mean), and ‘Length’ is the trimmed contig length (range and mean, bp). Ranks left as NA were never confidently resolved or were dropped as inconsistent with the contig phylum.

## Supplemental Figures

**Figure S1.**
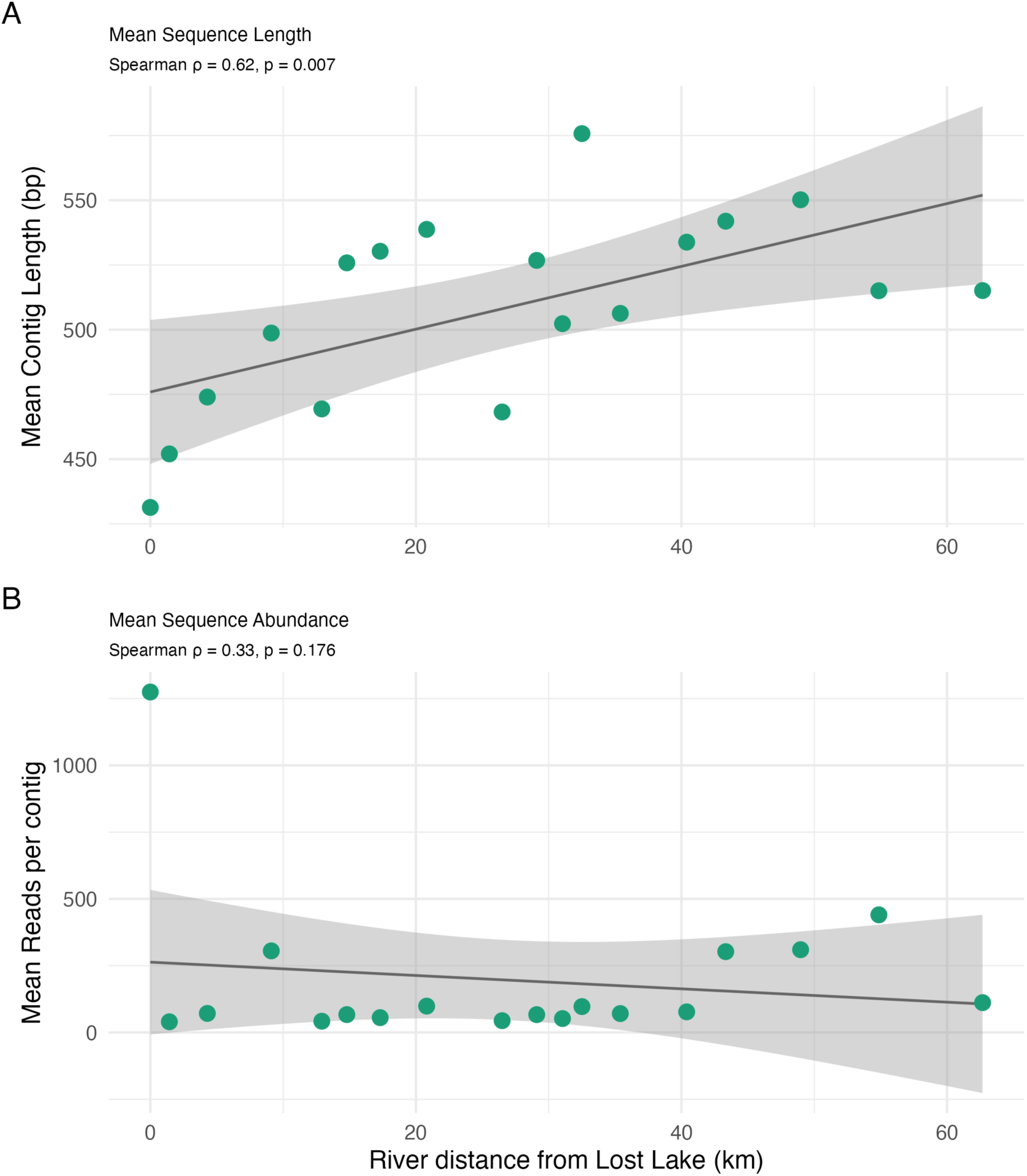
Invertebrate Sequence Characteristics vs River Distance. Mean contig length (top) and mean reads per contig (rel_abund, bottom) averaged per site for invertebrates, plotted against distance from the highest-elevation site. Spearman correlations: length rho = 0.62, p = 0.007; abundance rho = 0.33, p = 0.176.

**Figure S2.**
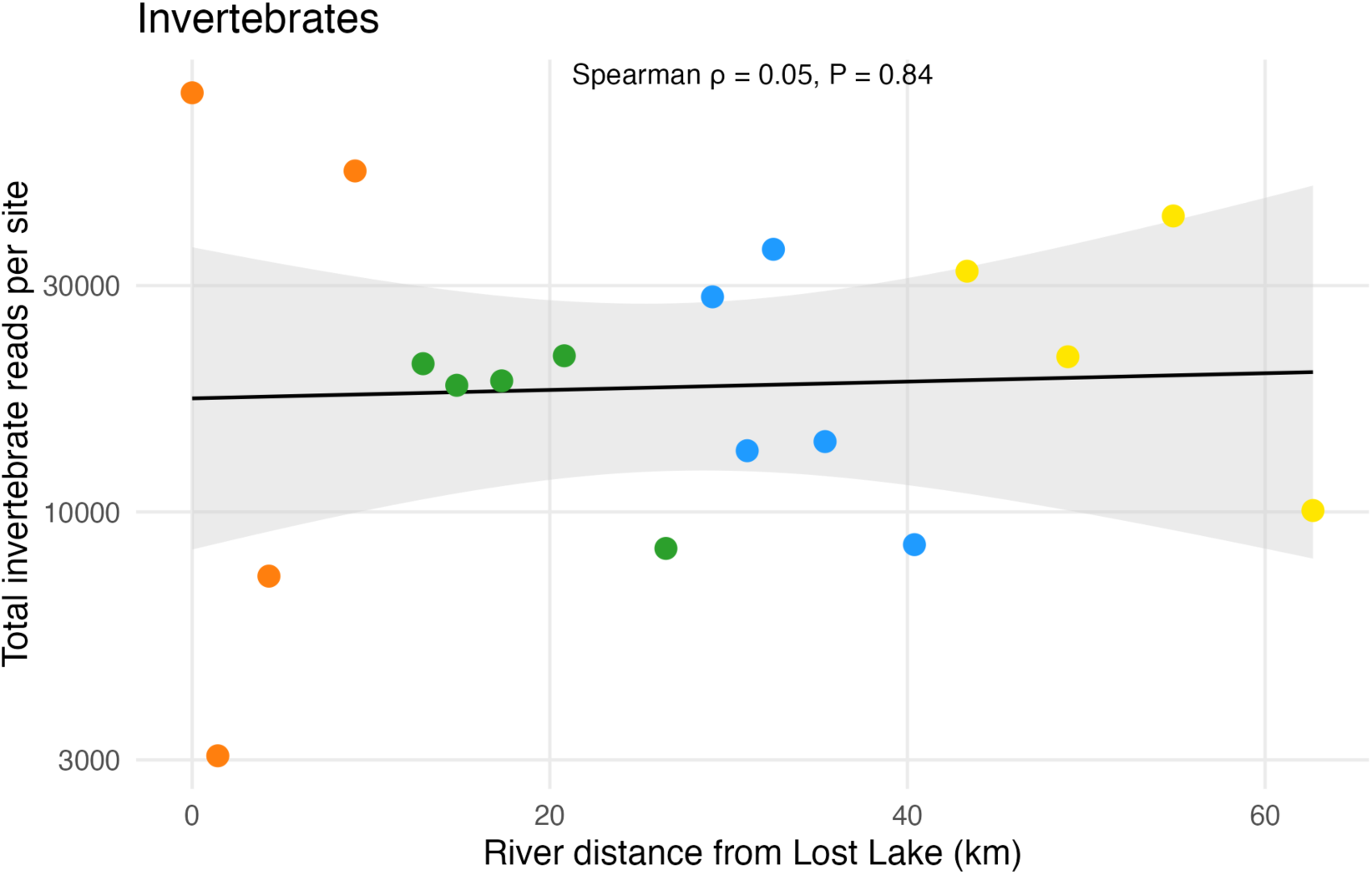
Read Depth vs River Distance. Site-level read depth—the sum of contig read counts across a site—plotted against river distance from Lost Lake. Invertebrate read depth shows no longitudinal trend (Spearman ρ = 0.05, P = 0.84; log10-linear R² = 0.002, P = 0.85)

**Figure S3.**
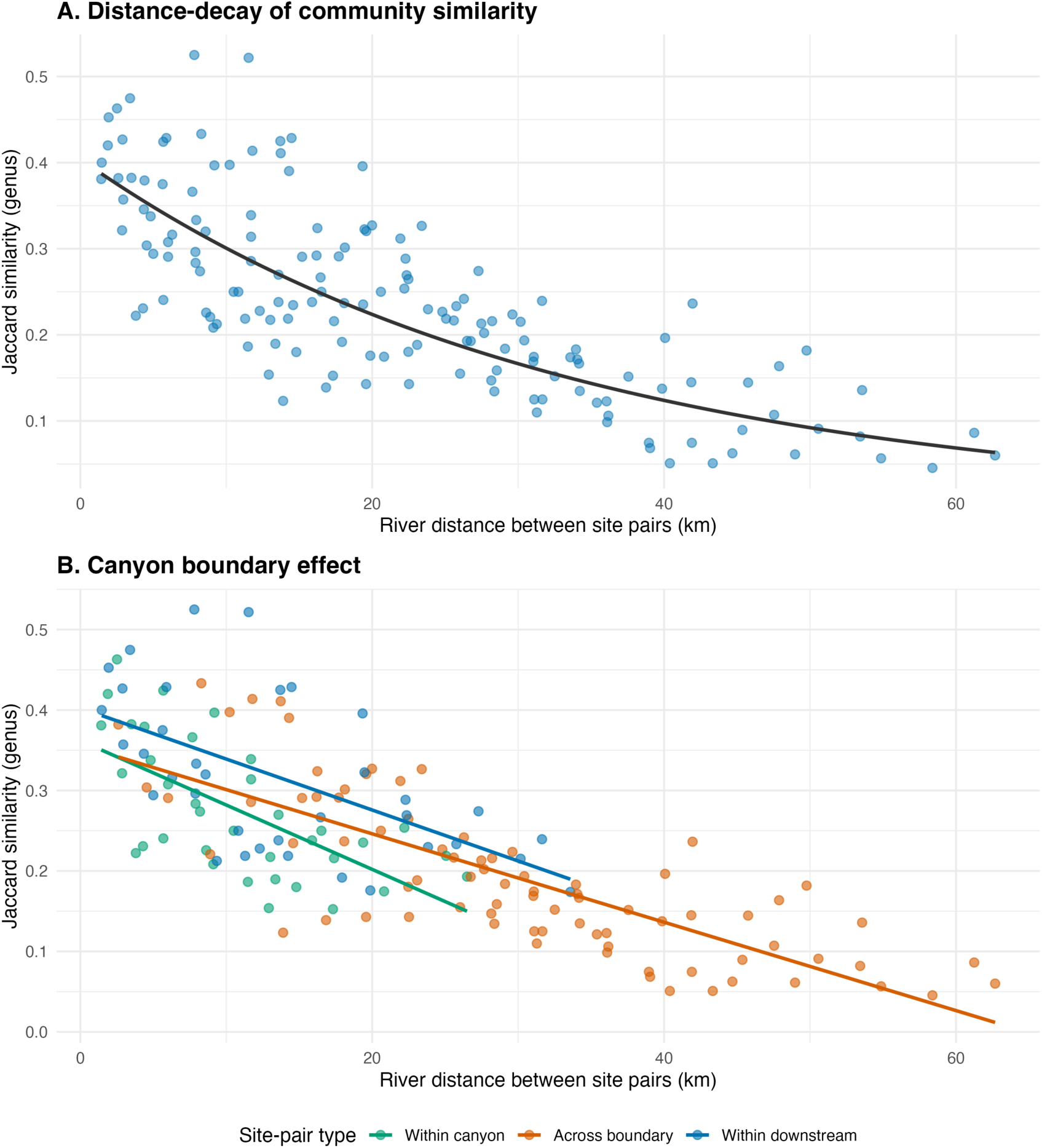
Distance-Decay of Invertebrate Community Similarity. (A) Pairwise genus-level Jaccard similarity between site pairs declines with river distance separating them (log-linear negative-exponential fit; Mantel r = 0.79, p = 0.001; similarity halves every ∼23 km). (B) Site pairs classified by zone: across-boundary pairs (one canyon site, one downstream site) are significantly less similar than within-zone pairs at the same distance (pair-type effect p = 0.002).

**Figure S4.**
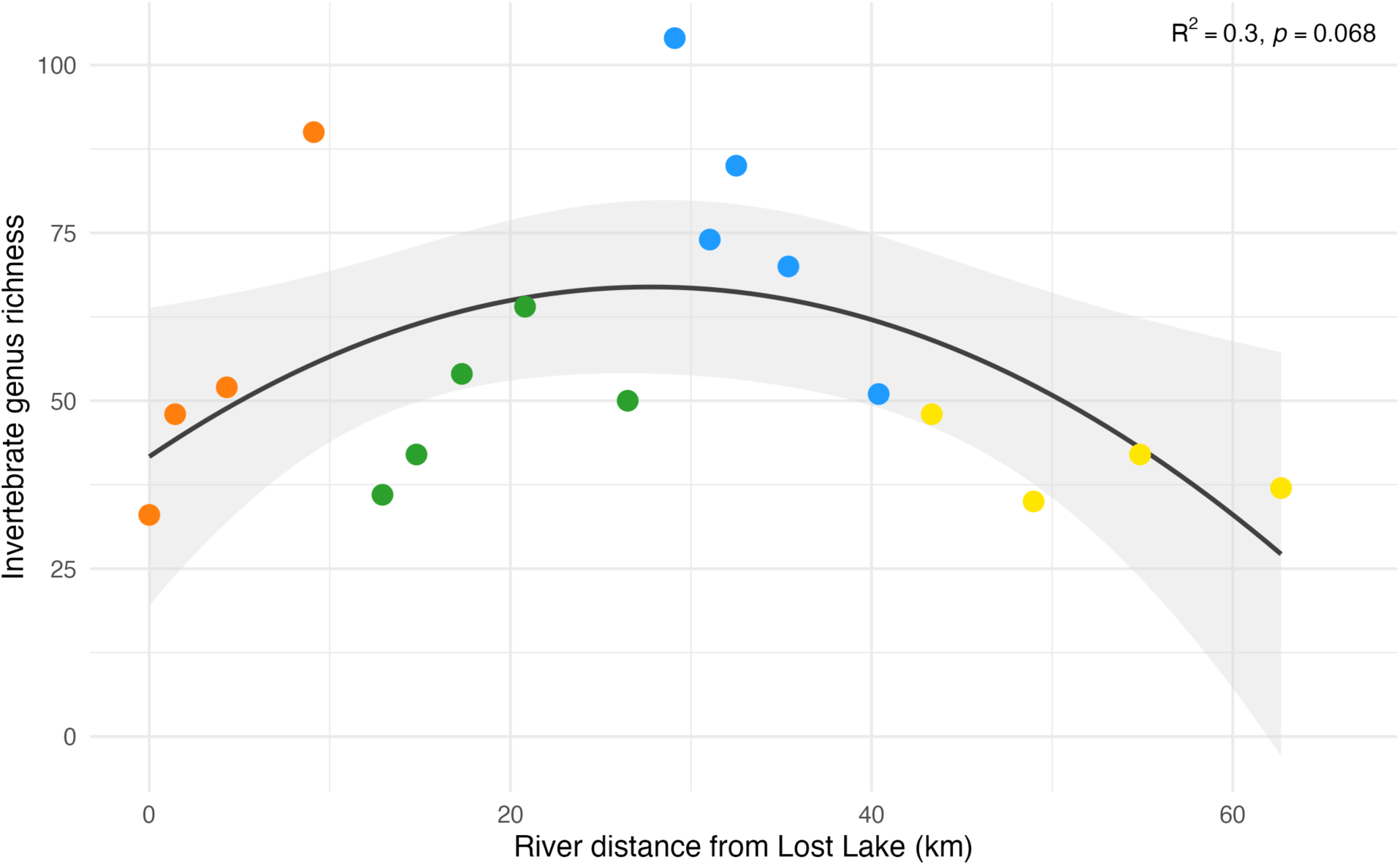
Taxonomic Richness Along the River Gradient. Invertebrate genus richness versus river distance from Lost Lake.

